# The Bacterial Sequential Markov Coalescent

**DOI:** 10.1101/090845

**Authors:** Nicola De Maio, Daniel J Wilson

## Abstract

Bacteria can exchange and acquire new genetic material from other organisms directly and via the environment. This process, known as bacterial recombination, has a strong impact on the evolution of bacteria, for example leading to the spread of antibiotic resistance across clades and species, and to the avoidance of clonal interference. Recombination hinders phylogenetic and transmission inference because it creates patterns of substitutions that are not consistent with the hypothesis of a single evolutionary tree (homoplasies). Bacterial recombination is typically modelled as statistically akin to the gene conversion process of eukaryotes, i.e., using the coalescent with gene conversion (CGC). However, this model can be very computationally demanding as it requires to account for the correlations of evolutionary histories of even distant loci. So, with the increasing popularity of whole genome sequencing, the need has emerged for a new and faster approach to model and simulate bacterial evolution at genomic scales. We present a new model that approximates the coalescent with gene conversion: the bacterial sequential Markov coalescent (BSMC). Our approach is based on a similar idea to the the sequential Markov coalescent (SMC), an approximation of the coalescent with recombination. However, bacterial recombination poses hurdles to a sequential Markov approximation, as it leads to strong correlations and linkage disequilibrium across very distant sites in the genome. Our BSMC overcomes these difficulties and shows both a considerable reduction in computational demand compared the exact CGC, and very similar patterns in the simulated data. We use the BSMC within an Approximate Bayesian Computation (ABC) inference scheme and show that we can correctly recover parameters simulated under the exact CGC, which further showcases the accuracy of our approximation. We also use this ABC approach to infer recombination rate, mutation rate, and recombination tract length from a whole genome alignment of *Bacillus cereus*. Lastly, we implemented our BSMC model within a new simulation software FastSimBac. In addition to the decreased computational demand compared to previous bacterial genome evolution simulators, FastSimBac also provides a much more general set of options for evolutionary scenarios, allowing population structure with migration, speciations, population size changes, and recombination hotspots. FastSimBac is available from https://bitbucket.org/nicofmay/fastsimbac and is distributed as open source under the terms of the GNU General Public Licence.

## Introduction

Whole-genome bacterial sequencing has rapidly replaced multilocus sequence typing as for population analyses of bacterial pathogens thanks to its fast and cost-effective provision of higher resolution genetic information (Didelot et al., 2012; Wilson, 2012). Methods using genomic data to infer epidemiological, phylogeographic, phylodynamic and evolutionary patterns are often hampered by recombination (e.g. Schierup and Hein, 2000; Posada and Crandall, 2002), and bacterial recombination is no exception (Hedge and Wilson, 2014). Recombination, in fact, causes different sites of the genome to have different inheritance histories. For these reasons, in recent years many methods have been proposed to measure, identify and account for bacterial recombination (e.g. Didelot and Falush, 2007; Marttinen et al., 2008; Tang et al., 2009; Didelot et al., 2010; Marttinen et al., 2012; Croucher et al., 2014; Didelot and Wilson, 2015). Among these, simulators of bacterial evolution (e.g. Didelot et al., 2009b; Mostowy et al., 2014; Brown et al., 2015) are used for parameter inference and hypothesis testing (e.g. Fearnhead et al., 2005; Fraser et al., 2005; Wilson et al., 2009; Ansari and Didelot, 2014) and for benchmarking (e.g. Falush et al., 2006; Didelot and Falush, 2007; Turner et al., 2007; Buckee et al., 2008; Marttinen et al., 2012; Hedge and Wilson, 2014).

Simulating bacterial evolution poses specific difficulties as the process of bacterial recombination is very different from that of other organisms. Eukaryotic recombination is predominantly modelled as a cross-over process, with recombination events breaking a chromosome into two parts with different ancestries (Figure 1). While it is possible to simulate eukaryotic evolution with recombination forward in time (Peng and Kimmel, 2005; Carvajal-Rodr Iguez, 2008; Hernandez, 2008; Arenas, 2013), coalescent-based (Kingman, 1982) backward in time models (Hudson, 1983; Griffiths and Marjoram, 1997; Wiuf and Hein, 1999) are usually more computationally efficient (e.g. Hudson, 2002; Arenas and Posada, 2007, 2010; Ewing and Hermisson, 2010; Excoffier and Foll, 2011). Yet, the coalescent with recombination itself may not be sufficiently fast when large genomic segments are considered (McVean and Cardin, 2005). One of the reasons is that the structure describing the evolutionary history of all positions (ancestral recombination graph, or ARG) seems to grow subexponentially with genome size and recombination rate (Wiuf and Hein, 1999). For this reason, a faster approximation to the coalescent with recombination, the sequential Markov coalescent (SMC, see McVean and Cardin, 2005; Marjoram and Wall, 2006) has been proposed. Similar to the sequential coalescent with recombination (Wiuf and Hein, 1999), the SMC starts by considering one evolutionary tree on the left end of the genome, and generates new trees affected by recombination as it moves towards the right end. However the SMC does not generate an ARG, but rather a sequence of local trees under the simplifying assumption that if the local tree for the current position is known, all previous local trees can be ignored in the next steps. This model has been further extended to include population history (Chen et al., 2009), increased accuracy (Wang et al., 2014), and increased computational efficiency (Staab et al., 2015).

Bacterial recombination is different from eukaryotic recombination (Smith et al., 1993, 2000), and is generally modelled as a gene conversion process, such that in a bacterial recombination event only a relatively small fragment of DNA is imported from a donor, whereas most of the genome is inherited clonally (Figure 1). This results in sites very distantly located in the genome to be very tightly linked genetically. In fact, a single genealogy, the clonal frame (Milkman and Bridges, 1990), represents the evolutionary history of all non-recombining sites, no matter how far they are from each other on the genome. For this reason, most methods used to describe and simulate eukaryotic recombination cannot be applied in bacteria. While bacterial evolution can also be simulated forward in time, backward in time coalescent methods are usually more efficient, and are generally based on the coalescent with gene conversion (CGC, see Wiuf and Hein, 2000, and Figure 2A). Recently, efficient methods implementing the CGC have been developed for simulating bacterial evolution (Didelot et al., 2009b; Brown et al., 2015). However, these approaches still struggle in simulating whole-genomes at high recombination rates (e.g. requiring up to hours to simulate a single bacterial genome-wide alignment with *ρ >*0.01, see Brown et al., 2015, and Results) because, similar to the coalescent with recombination, the CGC also generates large ARGs.

**Figure 1.**
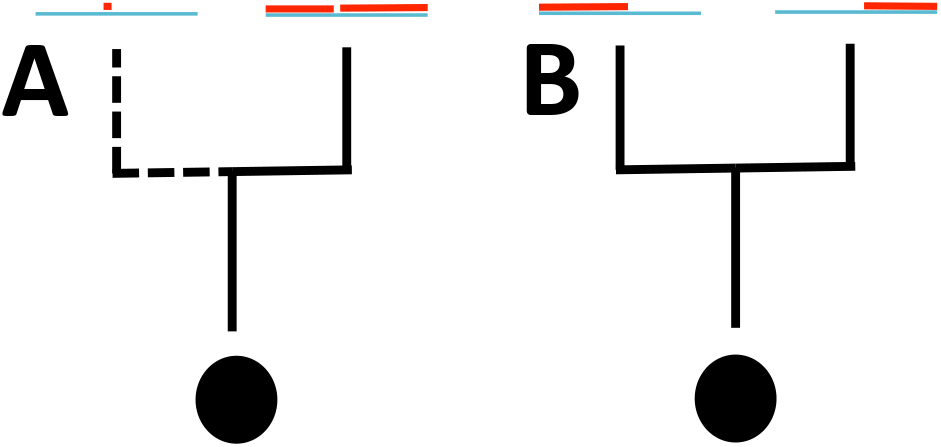
Graphical representation of bacterial and eukaryotic recombination models. Black circles represent samples, black lines are lineages (dashed if they represent bacterial recombination lineages). Blue segments represent the genome, and red segments represent the portion of the genome that is ancestral to the particular lineage. **A)** Gene conversion, or bacterial recombination: most of the genome is inherited from a single parent lineage, except for a short segment. **B)** Crossover event: all the genome on the left side of the crossover site is inherited from one parent, all the genome on the right side is inherited from the other parent.

Here, we present a new model of recombination (Figure 2B) that, inspired by the SMC, efficiently and accurately approximates bacterial recombination. We explicitly model the clonal frame, and simulate the coalescent and recombination processes along the genome conditional on the clonal frame, but “forgetting” recombination events that occur at distant positions. This approach differs by other approximations to the CGC (Didelot et al., 2010; Ansari and Didelot, 2014) as we can simulate entire genomes while allowing recombining lineages to coalesce with one another, and recombination events to split the ancestral material of recombinant lineages. Ignoring these complexities leads to biases when considering elevated recombination rates (Didelot et al., 2010), and by accounting for them we aim at specifying a model more adherent to the CGC. We call this model the bacterial sequential Markov coalescent (BSMC), which we implement within a new simulation software called FastSimBac. FastSimBac is faster than previous methods (between about one and two orders of magnitude for typical bacterial genome size and recombination rates). Also, by building on top of popular simulators ms (Hudson, 2002) and MaCS (Chen et al., 2009), our software can simulate more general evolutionary scenarios, allowing migration, speciation, demographic changes, recombination hotspots, and between-species recombination. We show that the BSMC can accurately approximate the exact CGC by inferring recombination parameters simulated under the CGC using Approximate Bayesian Computation (ABC) implementing BSMC simulations with FastSimBac. We also showcase its applicability by using it to infer recombination and mutation parameters via ABC from a whole genome alignment of *Bacillus cereus*.

**Figure 2.**
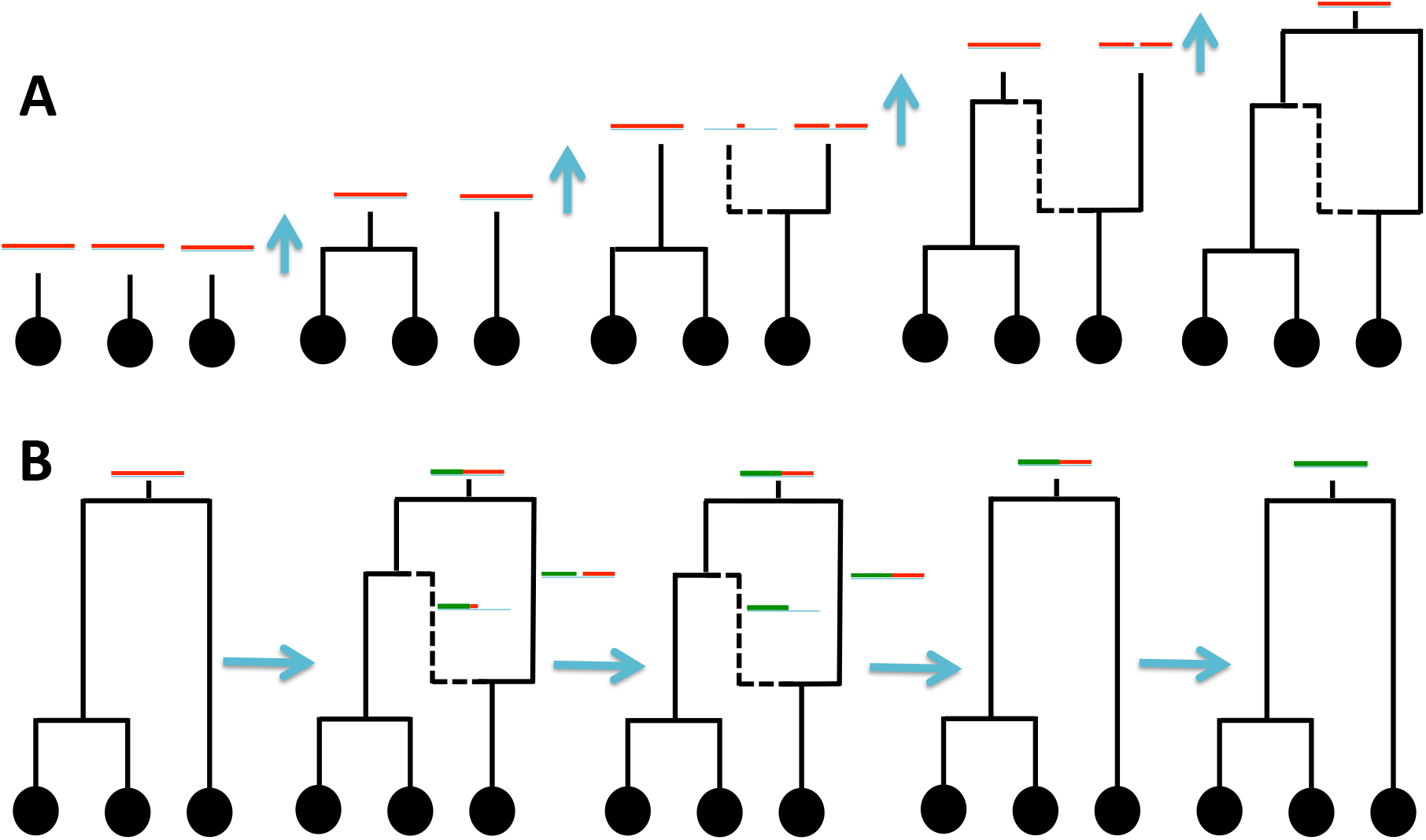
Graphical representation of the bacterial coalescent and BSMC models. Black circles represent samples, black lines are lineages (continuous if they belong to the clonal frame, dashed otherwise). Red segments represent, for each extant lineage, the portion of the genome that is ancestral to any sampled descendent of that lineage. Time is considered backward from bottom to top, and merges of lineages are coalescent events. **A)** Example simulation under the bacterial coalescent; recombinations and coalescent events are simulated backward in time starting from present, when one lineage per sample is given. **B)** Example of BSMC simulation: first a clonal frame is simulated; then, the process moves left to right across the genome (for simplicity the genome is assumed linear with left and right ends), and left portions of the genome are gradually forgotten (represented in green); recombination events are thus simulated at their start, and then forgotten at their end, but the clonal frame is never forgotten.

## Materials and Methods

### BSMC algorithm

We assume that a given set of parameters is priorly specified: *λ* is the mean length of a recombining segment, *G* is the total genome length, and *ρ* is the recombination rate. *λ* and *G* are considered in terms of base pairs, while *ρ*= 2*N_e_Gr* is the per-individual, per-generation, per-base pair gene conversion initiation rate *r* scaled by the effective population size *N_e_* and genome length *G*. Our BSMC algorithm, while inspired by the SMC (McVean and Cardin, 2005; Marjoram and Wall, 2006) in that it crosses the genome from left to right and discards previous local trees, also keeps tracks of and conditions on the clonal frame, and so has several important differences from the SMC. All lineages with ancestral material exclusively on the left of the currently considered genomic position *x_cur_* are forgotten (removed from the current local ARG *A*(*x_cur_*)), while all lineages with ancestral material on the right of *x_cur_* are stored in memory (included in *A*(*x_cur_*)). All lineages in *A*(*x_cur_*) are possible targets of new recombination events and coalescent events. Recombination events and coalescent events are instead not allowed on lineages that have been forgotten (not in *A*(*x_cur_*)). In order to decide which lineage is in *A*(*x_cur_*) and which needs to be removed, we record and update for each lineage *l* its ancestral material on the right of *x_cur_*: *al*(*x_cur_*). Updating the ancestral material of each lineage in *A*(*x_cur_*) after a new recombination has been added to *A*(*x_cur_*) is one of the most complex routines in our algorithm. One aim of the algorithm is to generate the sequence of local trees along the genome. For a given position *x_cur_*, the local tree (or marginal tree) *T*(*x_cur_*) is the coalescent tree describing the inheritance history of site *x_cur_*. *T*(*x_cur_*) can be obtained from *A*(*x_cur_*) by removing all branches that are not ancestral at *x_cur_*. While a simple graphical example of the algorithm is given in Figure S2, the list of BSMC algorithmic steps is the following:

1. *Initialisation: x_cur_*= 0 (current position, maximum is 1), and *T_cf_*(the clonal frame) is simulated under the coalescent without recombination. The initial local ARG *A*(*x_cur_*), and local tree *T*(*x_cur_*), are set to *T*(0) = *A*(0) = *T_cf_*. The ancestral material of every lineage *l* in *A*(0) is set to *al*(0) = [*x_cur_,*1] = [0,1], the whole genome. The list of recombination end points *E*(the right ends of recombination segments) is initialised as empty: *E*= ().
2. *Position of new event:* The distance until the next recombination initiation *x_new_* is drawn according to an exponential distribution 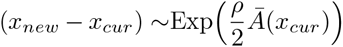, where 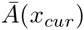 is the sum of all branch lengths in *A*(*x_cur_*), expressed in units of 2*N_e_* generations. If *_xnew_ > E*_0_, where *E*_0_ is the first (and smallest) element of the list *E* of recombination end points (if *E* is empty then *E*_0_ = *∞*), then *x_new_*= *E*_0_, *E*_0_ is removed from *E*, and the next event is a recombination termination, so go to step **4**. If *x_new_ ≥*1, and *E* is empty, terminate the algorithm. Otherwise the next event is a new recombination, so go to step **3**.
3. *New recombination event:* sample a lineage *l* randomly from *A*(*x_cur_*) proportionally to branch length. Then sample a time *t* uniformly along the time spanned by *l*. The new recombination happens at time *t* on branch *l*, and a new lineage *l’* is created, with its more recent end joining *l* at time *t*. A new coalescent time and coalescing lineage is sampled for *l’* conditional on *A*(*x_cur_*) (under the algorithm of Wiuf and Hein, 1999). The right end of the recombining interval *x_end_* is sampled from the distribution (*x_end_ − x_new_*) *∼ G_eom_*(*λ*)*/G*, where *Geom*(*λ*) is the geometric distribution with mean *λ*. If *x_end_ <*1, it is added to *E* in such a way to keep *E* sorted in increasing order. The new local ARG is defined *A*(*x_new_*) = *A*(*x_cur_*) *∪ l’* and ancestral material of all lineages in *A*(*x_new_*) is updated (ancestral material on the left of *x_new_* is deleted). All lineages with no ancestral material on the right side of *x_new_* are removed from *A*(*x_new_*). The new local tree *T*(*x_new_*) is defined from *A*(*x_new_*) and is printed to file. The current position is updated: *x_cur_*= *x_new_*. Go back to step **2**.
4. *Terminate a recombination event:* the ancestral material of all lineages in *A*(*x_new_*) is updated (ancestral material on the left of *x_new_* is deleted). All lineages with no ancestral material on the right side of *x_new_* are removed from *A*(*x_new_*). The new local tree *T*(*x_new_*) is defined from *A*(*x_new_*) and is printed to file. The current position is updated: *x_cur_*= *x_new_*. Go back to step **2**.

A large part of the complexity of the algorithm goes into the process of updating the ancestral material of lineages after a new recombination event is added to the local ARG. This step is described more in detail in the Supplement. Our algorithm and model differs from the approximation of the CGC used by (Didelot et al., 2010; Ansari and Didelot, 2014) in that, differently from them, we allow recombinant lineages to be affected by recombination, and to coalesce with each other if having overlapping ancestral material. To increase the realism of the model, we use the first positions simulated by the algorithm (generally 10*λ* bases) as burn-in, that is, they are simulated but not written to output or considered part of the genome length. While we simulate a linear genome, bacterial genomes are typically circular, so we assume that a genome start position has been arbitrarily chosen. The version of the algorithm above conveys the basics of the model of within-population recombination, and does not describe many additional events that we have included in our simulation software FastSimBac and that are described in the Supplement: mutations, migration, speciations, demographic changes, recombination hotspots and between-species recombination.

### Performance Testing

We simulated bacterial genome evolution under the coalescent with gene conversion using SimBac (Brown et al., 2015). We always simulated 50 contemporaneous samples. We performed simulations under four different recombination intensities:

*ρ*= 2*N_e_r*= 0.001,0.002,0.005,0.01, with *ρ* the population-scaled per generation per site recombination initiation rate. We also used four different genome sizes: *G*=1Mbp, 2Mbp, 5Mbp, 10Mbp. The mean recombination tract length was fixed to *λ*= 500. These values encompass a range of typical biologically relevant scenarios for bacteria (Vos and Didelot, 2009; Didelot and Maiden, 2010). We simulated 10 replicates for each combination of parameters, and for each replicate the collection of local trees, and the clonal frame, were stored. Sequence data was generated using the local trees and SeqGen (Rambaut and Grassly, 1997) under an HKY85 model (Hasegawa et al., 1985) with transition/transversion rate ratio *κ*= 3. Some of the parameter combinations were too computationally demanding for SimBac: (*ρ*= 0.005*, G*=10Mbp), (*ρ*= 0.01*, G*=5Mbp), (*ρ*= 0.01*, G*=10Mbp). For all the replicates for which we could run SimBac, we used the clonal frame simulated by SimBac as an input for our new software FastSimBac. In fact, the clonal frame is a major source of variation in sequence patterns between simulations (Ansari and Didelot, 2014). By using the same clonal frames in the two methods we expect less variance in the difference of summary statistics between the two methods; in particular, we eliminate the variance associated with the clonal frame, and this allows us to perform fewer simulations to compare the methods. For all scenarios in which we could not run SimBac, the clonal frame was generated randomly within FastSimBac. Again, we generated local trees in FastSimBac and used these to generate alignments in SeqGen as before.

### Approximate Bayesian Computation Inference

We performed Approximate Bayesian Computation (ABC) inference with the local-linear regression approach (Beaumont et al., 2002) as implemented in the R package abc (Csilléry et al., 2012). To test the performance of an ABC scheme based on our BSMC model, we used it with FastSimBac simulations to infer parameters from datasets simulated under the CGC using SimBac. We used a uniform prior distribution over [0,0.005] for the recombination rate *ρ*, and over [10,1000] for the mean length *λ* of recombining intervals. The same priors were used for simulating datasets and for performing inference. The aim of the ABC analyses was to infer these *ρ* and *λ*. For simplicity, the clonal frame simulated in SimBac was assumed to be known (see also Ansari and Didelot, 2014; Hedge and Wilson, 2014), as was the mutation rate *θ*= 0.005. The genome size was fixed to 1Mbp, and the number of samples to 20. For each true data set simulated with SimBac, we simulated 10,000 approximate datasets under the BSMC in FastSimBac. Only 1% of the simulations in FastSimBac was retained for parameter inference (the 1% with closest summary statistics to the true dataset, see Beaumont et al., 2002). We used two summary statistics: the proportion of incompatible sites (G4) between neighbouring SNPs, and the G4 between SNPs at least 20kbp away. More precisely, we considered the simulated alignment starting from the left end of the genome, and, for the first summary statistic, for each SNP we selected the first SNP occurring on its right; for the second summary statistic for each SNP we selected the first SNP on its right at least 20kbp away. The idea is that G4 (and linkage disequilibrium) at very short distances (*≪ λ*) will mostly depend only on the recombination rate *ρ*, while G4 on long distances (*≫ λ*) will mostly depend on the product *ρλ*, so that these two summary statistics together will give sufficient information to estimate *ρ* and *λ*.

We also performed Approximate Bayesian Computation (ABC) inference on a real *Bacillus cereus* genome alignment (Didelot et al., 2010; Ansari and Didelot, 2014) with the ABC-MCMC scheme (Marjoram et al., 2003). We used uniform prior distributions on [0.0,0.25] for *ρ*, on [1,10000] for *λ*, and on [0.01,0.2] for *θ*(the per-bp per-individual per-generation mutation rate scaled by 2*Ne*). These are the 3 parameters that we attempt to infer. We simulated entire genome alignments of 13 samples and 5240935 bp, as for the real dataset. We use 7 summary statistics: number of polymorphic sites (real value 629942); G4 (proportion of SNP pairs that are not consistent, breaking the 4-gamete rule) for consecutive SNPs (real value 0.167) and for SNPs at least 2kbp away (real value 0.297); mean linkage disequilibrium (LD, measured as 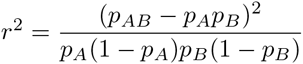 where *p_A_* is the frequency of allele A in the first SNP, *p_B_* the frequency of B in the second SNP, and *p_AB_* the frequency of the AB haplotype) for consecutive SNPs (real value 0.396) and for SNPs at least 2kbp away (real value 0.274); and mean number of haplotypes (considering a certain number of SNPs at the time) for pairs of consecutive SNPs (real value 3.003) and for groups of 4 SNPs made of 2 pairs of consecutive SNPs, the two pairs being at a distance of at least 2kbp. While the number of SNPs is informative of the mutation rate, the three summary statistics at short range are informative of the recombination initiation rate, while the three summary statistics at long range are informative of the product *ρλ*. Number of SNPs, G4 and *r*2 were also used as summary statistics by (Ansari and Didelot, 2014). The fact that we are able to generate entire genomes (instead of SNP pairs as Ansari and Didelot, 2014) allows us also to include summary statistics on groups of SNPs, such as numbers of haplotypes. For simplicity we fixed the clonal frame to the one estimated and used by (Didelot et al., 2010; Ansari and Didelot, 2014). However, we also correct for the branch lengths estimation error caused by recombination. In fact, with increasing recombination, all genetic distances between samples converge to a unique value. We discuss this bias and our corresponding correction in the Supplement. Lastly, in an attempt to further increase the realism of our model, we account for invariable sites. In fact, a large proportion of the sites is polymorphic (about 1 bp every 6 after removing sites with limited coverage) and a large proportion of the genome is expected to be coding; so, in principle, one would expect many homoplasies (sites patterns not consistent with the clonal frame and the infinite sites assumption) to occur just due to multiple mutations at one site, and not necessarily involve recombinations. Using back of the envelop calculations (see Supplement) we estimated about half of the genome to be invariant (48.44% of sites) and a transition-transversion ratio of about 5.21. We used these estimates as fixed values within an HKY (Hasegawa et al., 1985) substitution model with invariant sites, instead of the basic JC model (Jukes and Cantor, 1969) implemented in our basic inference and in (Ansari and Didelot, 2014). This model, together with the local trees simulated by FastSimBac, was used in SeqGen to simulate the alignment from which summary statistics were extracted at each step of the ABC-MCMC. Each run consisted of 10000 ABC-MCMC steps, of which 1000 were used as burn-in.

## Results and Discussion

### Computational efficiency of BSMC

Thanks to our BSMC approximation that simplifies the coalescent with gene conversion (CGC) by considering many small local ARGs, instead of a unique, large, global ARG, FastSimBac shows great computational improvement in simulating typical bacterial genome evolution. Compared to the currently most efficient software to the best of our knowledge, SimBac, FastSimBac speed improvements range from about one order of magnitude for low recombination rate (*ρ*= 0.001) and genome size(106bp), to two orders of magnitude for more elevated recombination rate (*ρ*= 0.01) and genome size (107bp), as shown in Figure 3. Also, FastSimBac allows simulation of scenarios with both high recombination rate and genome size which are currently out of reach of other methods, due to excessive requirements in time and RAM. In fact, we see that the performance of FastSimBac relative to the exact coalescent with gene conversion improves as we increase either genome size or recombination rate (Figure 3). As expected, the running time of FastSimBac appears linear with genome size, while this is not true for SimBac. Another benefit of FastSimBac is that, by avoiding the generation of a global ARG, it has small RAM usage, which allows it to efficiently run in parallel on multiple cores.

### Accuracy of BSMC

Next, we compare the simulated patterns of genetic variation and local tree features between the exact CGC simulated under SimBac, and the BSMC simulated with FastSimBac. Looking at linkage disequilibrium (LD, measured as *r*^2^) and site pair incompatibility (or four-gamete test, G4), we notice that, as expected, LD decreases and G4 increases considerably with increasing recombination rate (Figure 4). There is a lot of variation across different replicates in mean LD, but this is also expected as each replicate has a distinct clonal frame, and the clonal frame influences site patterns of the whole genome. LD and G4 at 1kbp are already very close to that of longer distances, suggesting that a distance of 2*λ* is sufficient to reach nearly as much LD as any arbitrary distance. Most importantly, we notice that values simulated under the BSMC mimic very closely those simulated under the exact CGC, suggesting that indeed, even at high recombination rates and short distances, the BSMC is a very accurate approximation (Figure 4). Similar results are also observed at different genome sizes (Figure S3).

**Figure 3.**
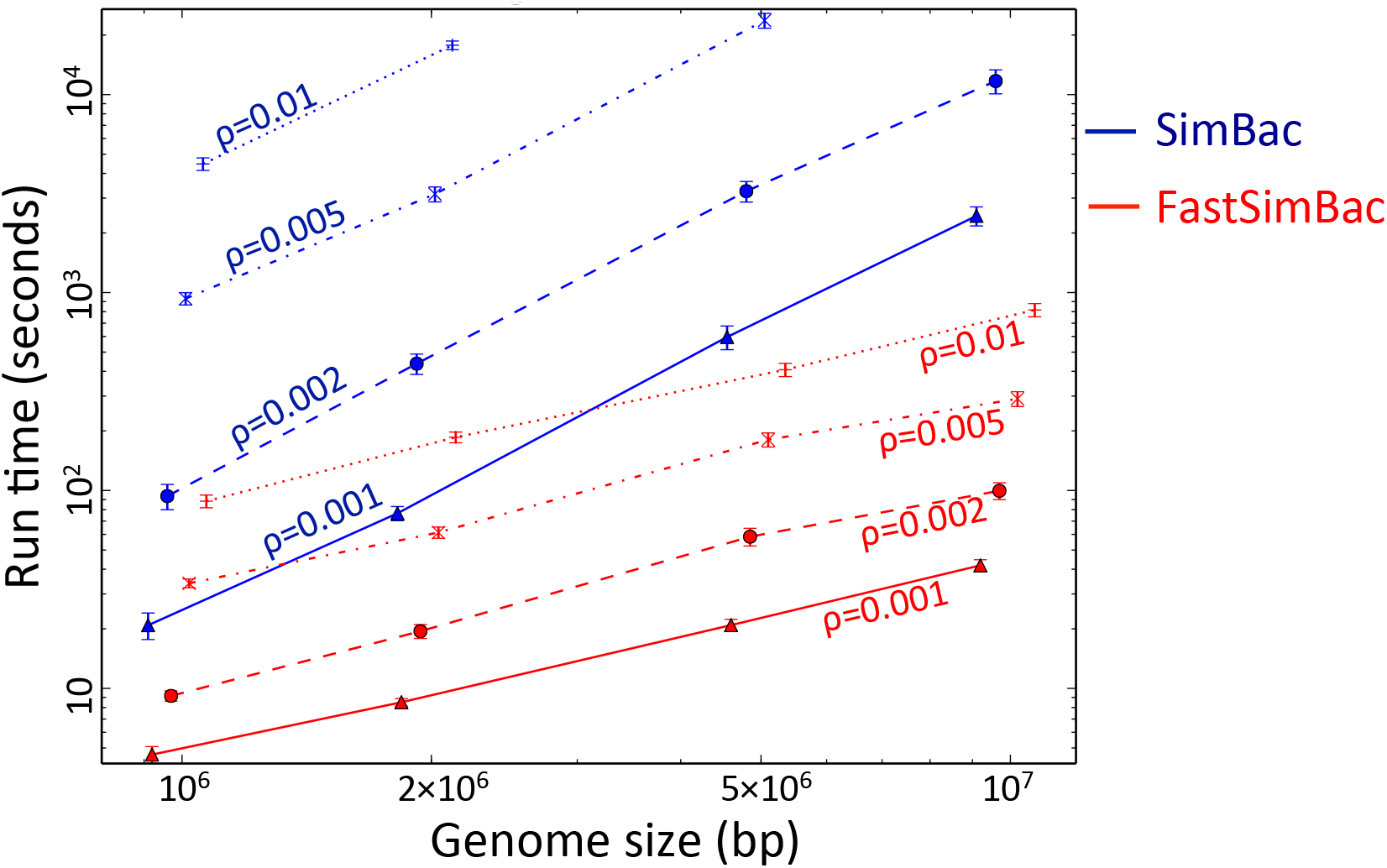
Comparison of computational demand between the BSMC and the coalescent with gene conversion (CGC). The bacterial sequential Markov coalescent implemented in FastSimBac is faster than the exact CGC implemented in SimBac. On the Y axis is the per replicate running time to generate local trees (in seconds and on a log scale), on the X axis the genome size (in bp and on a log scale). Red lines refer to FastSimBac, blue lines to SimBac. Each point is the mean over 10 replicates, and bars are standard errors of the mean. SimBac was not run for the combinations of highest recombination rates and genome sizes due to time limitations.

**Figure 4.**
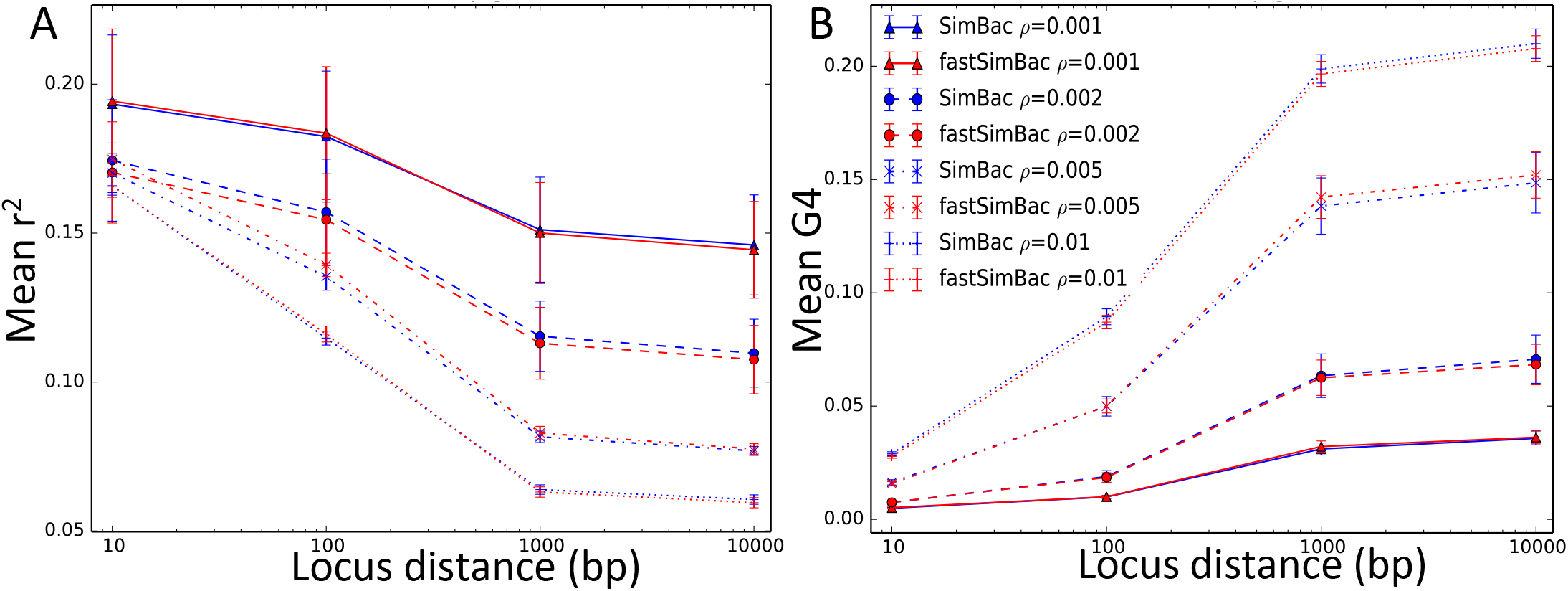
Comparison of linkage disequilibrium and site incompatibility between the BSMC and the CGC. The BSMC has patterns of linkage disequilibrium (LD, measure as *r*^2^) and site incompatibility (G4) very similar to the bacterial coalescent. On the X axis is the distance between SNPs in bp at which LD and G4 are measured. *r*^2^ is calculated as 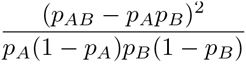, and G4 (the four-gamete test) is 1 if a SNP pair is incompatible and 0 otherwise. For each distance *d*, and for any SNP *x*, LD and G4 were calculated between *x* and the first SNP at least *d* bp to the right of *x*. Red lines refer to FastSimBac, blue lines to SimBac, and different point and line styles refer to different recombination rates (see legend). Genome length was fixed at 1Mbp. Each point is the mean over 20 replicates, and bars are standard errors of the mean. **A)** Mean LD. **B)** Mean G4.

Additionally, looking at the number of haplotypes present in non-overlapping windows of 10 SNPs, we observe an expected increases with recombination rate (Figure 5A). More importantly, the BSMC again very closely mimics the exact CGC. The genomic variation in number of haplotypes (Figure S4A) is very slightly underestimated, probably because long-range correlations in local trees (after conditioning on the clonal frame) are ignored in the BSMC, while present in the CGC. The mean pairwise genetic distances between samples appears not affected by recombination and by the model used for simulations (Figure 5B), but recombination does affect the variance of genetic distances over sample pairs (Figure S4B) because it tends to break down the relatedness of samples. Again, both patterns in the CGC are very closely approximated by the BSMC. Looking at mean local tree height (Figure 5C) and mean local tree total branch length (Figure 5D) we see that these are highly variable dependent on the simulated clonal frame, but are not considerably affected by the simulation parameters. Again, BSMC and CGC values are very close.

### BSMC-based ABC inference

We investigated the accuracy and applicability of the BSMC approximation by performing ABC inference of parameters. First, we reconstructed parameters simulated under the exact CGC. We use summary statistics based incompatibilities indicative of recombination between pairs of sites (G4). Despite the fact that the exact CGC was used to create the original datasets, while our BSMC was used for the ABC, inference was accurate. 95% posterior confidence intervals for *ρ* and *λ*(respectively the population-scaled recombination rate and the mean length of recombining intervals) contain the simulated values in both our replicates (Figure 6 and S5). This supports the idea that sequential Markov approximations of the CGC can be used for accurately inferring bacterial evolutionary parameters.

As an additional example of the applicability of our model and software, we used an ABC-MCMC approach (Marjoram et al., 2003) to infer *ρ*, *λ*, and the scaled mutation rate *θ* for the *Bacillus cereus* bacterial group. Bacteria of the *B. cereus* group mostly live in the soil, feeding on dead organic matter, but they can occasionally infect humans and cause a range of diseases, from food poisoning up to deadly anthrax (Arnesen et al., 2008). Disagreement has been found between *B. cereus* species designation and MLST clade structure and population history, probably due to the contribution of plasmids and genetic recombination to the bacterial phenotype (Priest et al., 2004; Sorokin et al., 2006; Didelot et al., 2009a; Zwick et al., 2012). Furthermore, analyses of MLST data showed discordant results regarding the prevalence of recombination in *B. cereus* relative to mutation, with estimates ranging from *ρ/θ ≈*0.05 (Hanage et al., 2006), to *ρ/θ ≈*0.2 (Didelot et al., 2009a), to *ρ/θ ≈*0.3 (Didelot and Falush, 2007), up to *ρ/θ ≈*2 (Pérez-Losada et al., 2006), leading to present uncertainty regarding the contribution of recombination to the *B. cereus* evolution. Improving our understanding of recombination in *B. cereus* would help us recognise the effect of homologous recombination on epidemiological inference and species delimitation (Didelot and Maiden, 2010), and predict the acquisition and spread of infectivity and resistance factors (Perron et al., 2011). With this respect, genome-wide data from multiple strains provide a greater opportunity to study recombination in detail, and here we consider the genome alignment described in Didelot et al. (2010) and Ansari and Didelot (2014), and comprising 13 genomes from the *B. cereus* group. Didelot et al. (2010) performed MCMC inference on this dataset using an approximate coalescent model with bacterial recombination (the ClonalOrigin model) that did not allow recombinant lineages to be affected by further recombination, or recombinant lineages to coalesce with each other. They inferred a mean recombination tract length of *λ*= 171bp with interquartile range [168,175], and *ρ/θ*= 0.21 with interquartile range [0.20,0.23]. Ansari and Didelot (2014) used again a model similar to the ClonalOrigin one within an ABC-MCMC approach, and accounted for the propensity of lineages to recombine with more closely related lineages than with distantly related ones. They inferred *ρ*= 0.077 with confidence interval *CI_ρ_*= [0.036,0.127], *λ*= 152bp with *CI_λ_*= [74,279], and *θ*= 0.0528 with *CI_θ_*= [0.0437,0.0640]. The ClonalOrigin model used by these methods approximates the coalescent with gene conversion, but in a less adherent way than the BSMC; in fact, their model leads to overestimation of the recombination rate *ρ* at elevated recombination and mutation rates which are relevant in this scenario (Didelot et al., 2010). Our BSMC-based ABC-MCMC approach instead allows recombining lineages to coalesce with one another, and recombination events to split the ancestral material of recombinant lineages. Furthermore, differently from these previous methods we account for differences in transition and transversion rates, for invariant sites, and for biases in tree branch length estimation (see Materials and Methods and Supplement)

**Figure 5.**
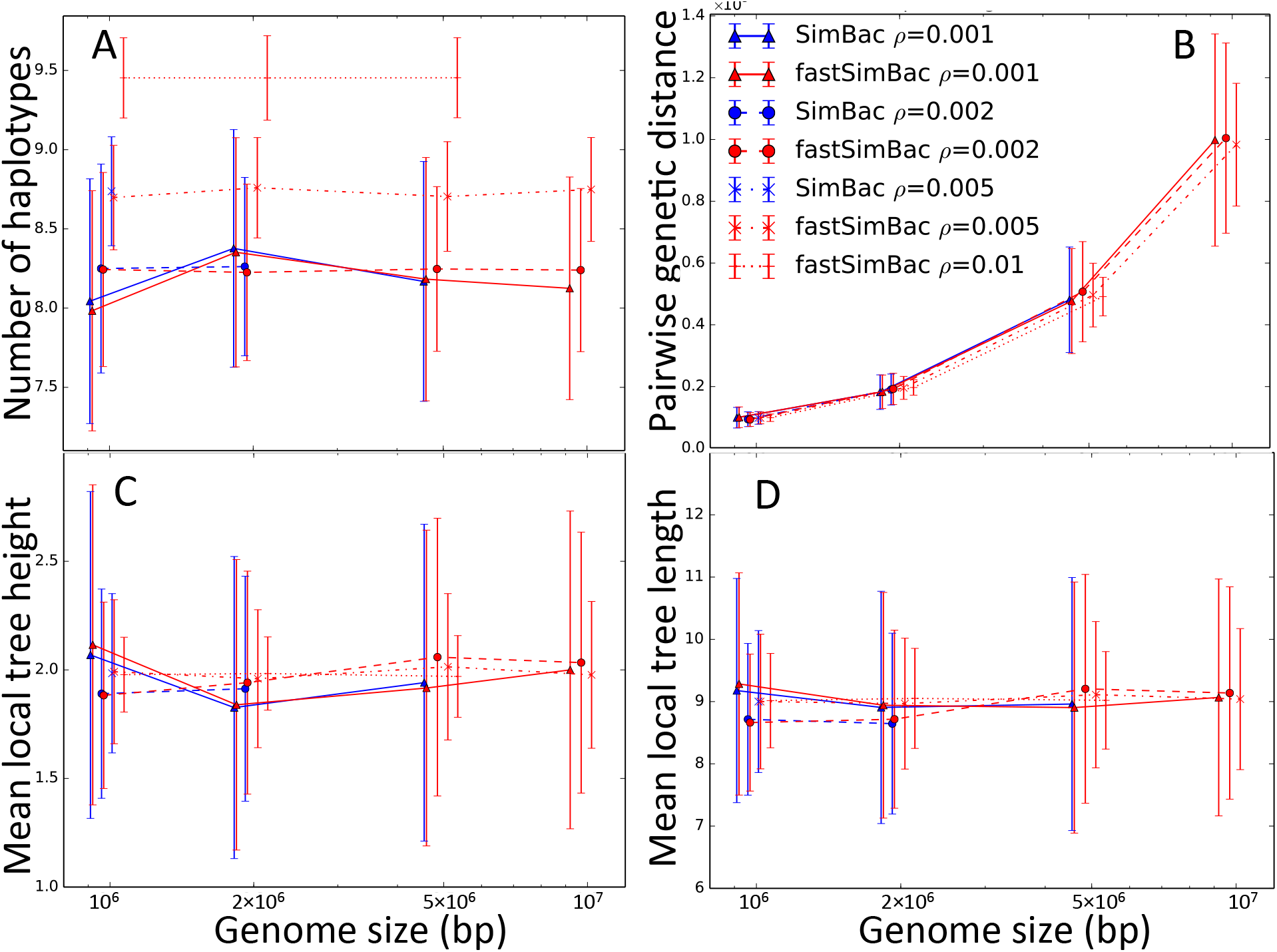
Comparison of several simulated patterns between the BSMC and the exact CGC. Bacterial evolution simulated under the BSMC gives very similar results to the exact CGC. **A)** Genome-wide mean number of simulated haplotypes over non-overlapping sliding windows of 10 SNPs; **B)** Mean over all sample pairs of whole-genome genetic differences; **C)** Mean local tree height; **D)** Mean local tree length (sum of all branch lengths). On the X axis is genome size in bp and on log scale. Red lines refer to FastSimBac, blue lines to SimBac, and different line and dot styles refer to different recombination rates (see legend). Each point is the mean over 50 replicates, and bars are standard deviations. SimBac and FastSimBac were not run for the highest recombination rates and genome sizes due to time and memory limitations.

**Figure 6.**
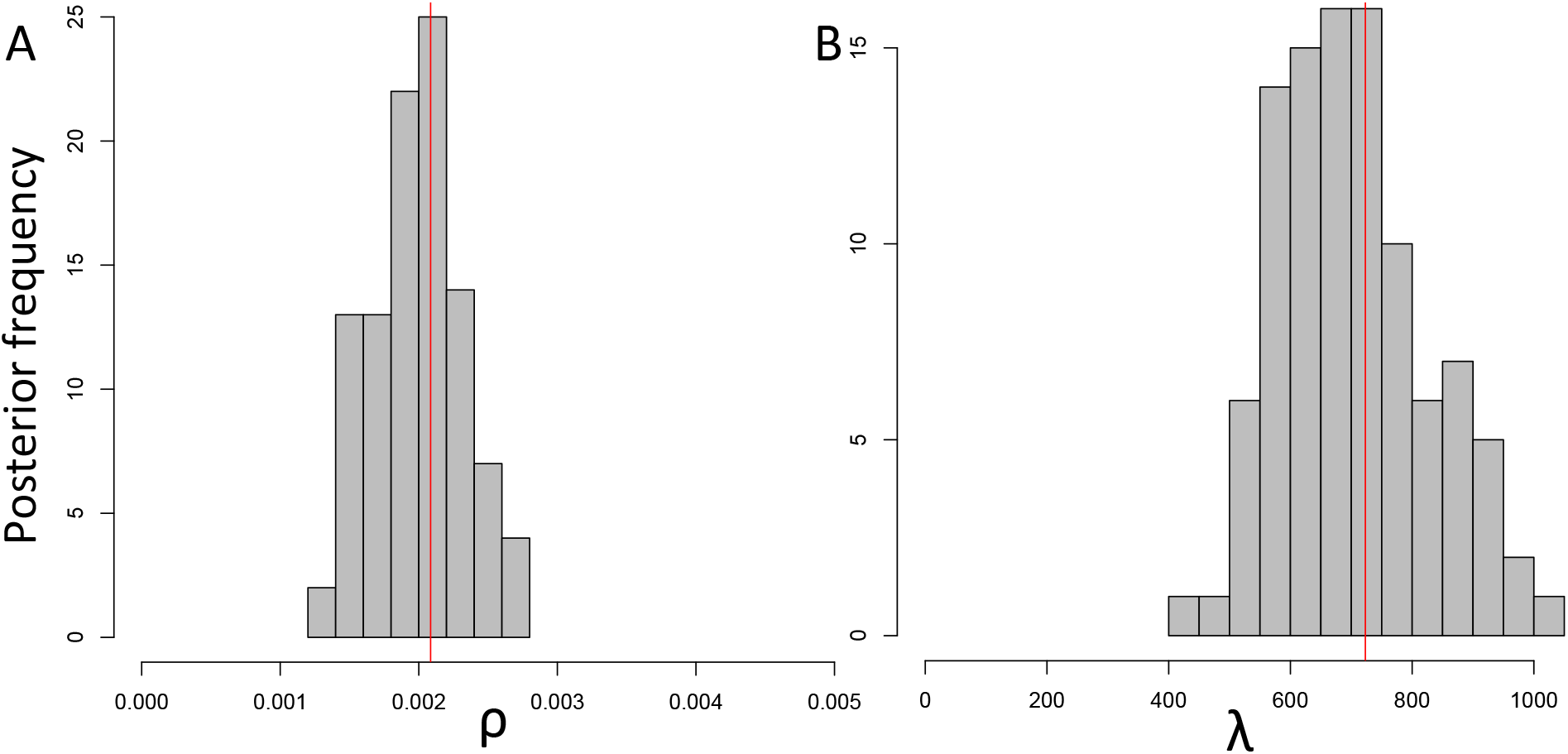
Accurate inference of recombination parameters with the BSMC-based ABC. Recombination parameters simulated under the exact CGC (red vertical lines) where reconstructed using simulations under the BSMC within an ABC inference scheme. Inference from another independent ABC run is shown in Figure S5. **A**) Posterior distribution of *ρ*. **B)** Posterior distribution of *λ*.

**Figure 7.**
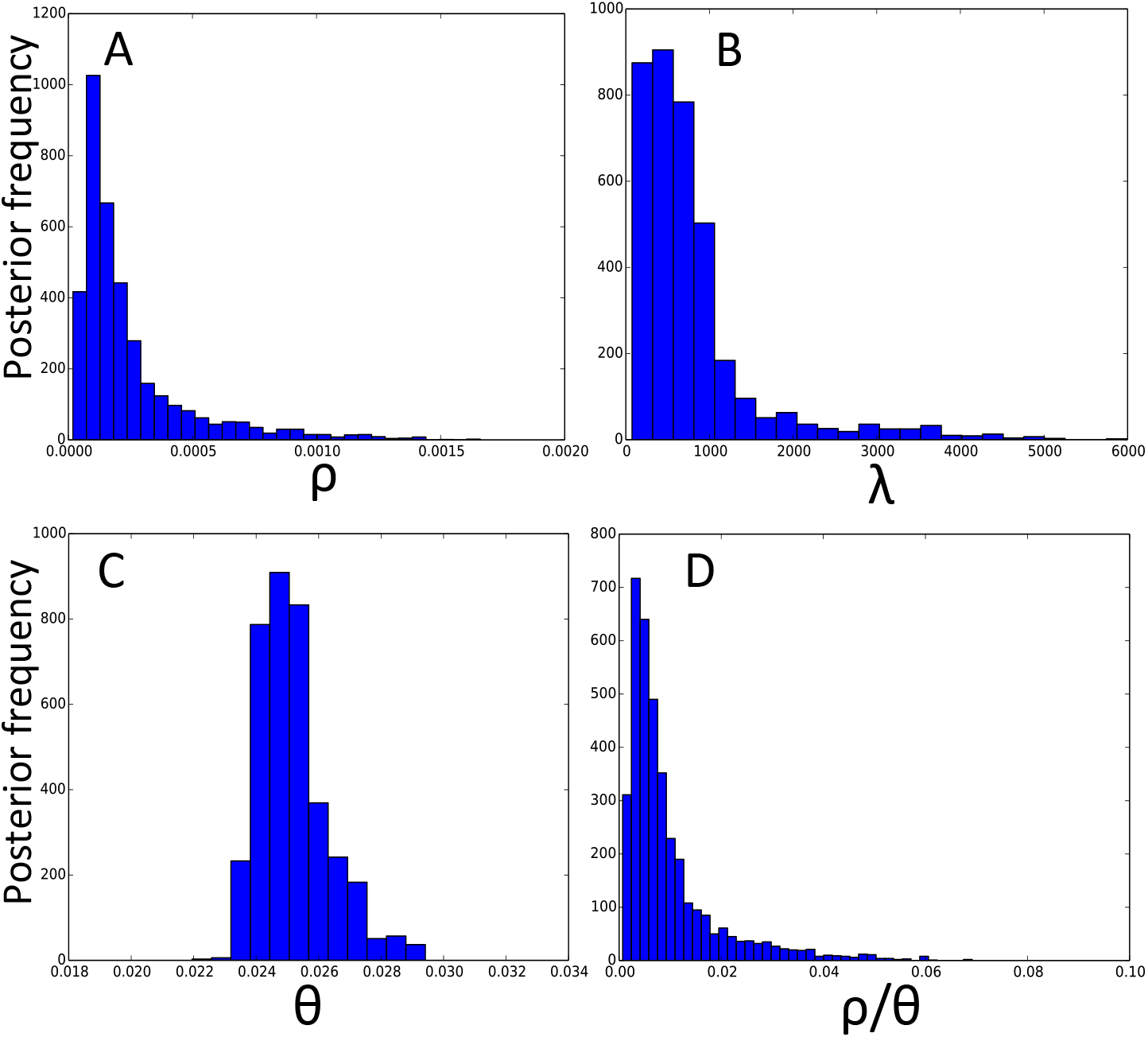
Posterior distributions of parameters for genome-wide evolution of *B. cereus*. We inferred BSMC parameters using an ABC-MCMC inference scheme. A) Posterior distribution of *ρ*. **B)** Posterior distribution of *λ*. **C)** Posterior distribution of *θ*. **D)** Posterior distribution of *ρ/θ*.

With our BSMC-based approach, we inferred higher mean recombination tract length *λ*(median 592bp and interquartile range [336,885]) than previous estimates (52bp and 171bp from Didelot et al. (2010) and Ansari and Didelot (2014) respectively); This estimate is closer to values inferred from genome-wide likelihood-based analyses in *Clostridium difficile*(Didelot and Wilson, 2015). We also inferred a considerably lower contribution of recombination relative to mutation (*ρ/θ*, median 0.0065 and interquartile range [0.004,0.011]) than previous genome-wide studies (0.21 and *≈*1.46 from Didelot et al. (2010) and Ansari and Didelot (2014) respectively); this means that recombination has a much lower contribution to evolution in *B. cereus* than previously thought, and that in fact these bacteria are considerably clonal, although, due to variation in recombination rates between *B. cereus* clades, our results do of course not apply to all species within the *B. cereus* group (Sorokin et al., 2006). These results were confirmed by an additional independent estimation run (Figure S7), and can be explained by the fact that we account for invariant sites and for different transition and transversion rates. In fact, invariant sites and high transition/transversion rate ratio cause more homoplasies than expected under an homogeneous substitution rate; these homoplasies, if unaccounted for, can be interpreted as short recombinant fragments, biasing downward estimates of *λ*, and upward estimates of *ρ/θ*. Supporting our interpretation, when we ran our method without accounting for invariant sites we estimated lower *λ* and higher *ρ/θ*(Figure S8). Another factor that can explain our larger *λ* estimate is that in our BSMC model we allow recombination events to interfere with each other, breaking recombinant segments into smaller pieces as expected in the CGC, and this process, if unaccounted for, could lead to a downward bias in the estimation of *λ*.

We also found correlation between *ρ* and *λ*, suggesting that while the total impact of recombination *ρ * λ* is easier to estimate, identifying the two individual parameters is harder. We found no correlation instead between other pairs of parameters (Figure S6 A-C, see also Ansari and Didelot, 2014). While our ABC-MCMC seems to capture well the complexity of real data for 5 out of 7 summary statistics, for two of them (G4 at large distances and *r*2 at short distances) there seem to be discrepancies (Figure S6 D-J and Figure S7 E-K). This might suggest the existence of some complexities that we did not account for in our model, for example the larger rate of recombination between closely related lineages (see Ansari and Didelot, 2014), variable recombination rate between *B. cereus* clades (Sorokin et al., 2006), prevalent non-homologous recombination (Didelot and Maiden, 2010), population structure (such as due to niche adaptation Sorokin et al., 2006), recombination with other bacterial groups, variable selective pressure and mutation rate, and alignment errors.

In conclusion, the BSMC offers not only a very computationally convenient approximation to the CGC, but also an accurate one. Our implementation of the BSMC model in the simulation software FastSimBac allows faster simulations, and therefore parameter inference, of bacterial genome evolution, and under a broader range of parameter values. FastSimBac also allows specification of the clonal frame upon which to condition simulations, which can grant simulations a closer fit to particular phylogenies reconstructed from real datasets. But more importantly, by virtue of building on top of the popular simulators ms (Hudson, 2002) and MaCS (Chen et al., 2009), our software includes many evolutionary scenario options that have been included in previous eukaryotic coalescent simulators (Hudson, 2002; Chen et al., 2009) but have remained precluded from bacterial coalescent simulations, such as population structure and migration, speciation histories, changes in population sizes, and recombination hotspots. FastSimBac is available as open source from https://bitbucket.org/nicofmay/fastsimbac. Applications of our model and software are not necessarily restricted to simulations, but, as we have shown, also include inference of recombination rate and other parameters of bacterial evolution. Our simulations suggest that our approach gives results very close to those obtained with the exact CGC, but at considerably reduced computational cost. Our analysis of recombination in the *B. cereus* group showcases the applicability of our method for inference from genome-wide alignments. Possible further applications of our model include the study and inference of recombination patterns and events, or the estimation of the clonal frame while accounting for recombination. We thus believe that the BSMC and FastSimBac will provide very useful for both benchmarking and for statistical inference on bacterial whole genome sequence data.

## Acknowledgements

We are grateful to the creators of MaCS, on which the FastSimBac code is partly based. We also thank Xavier Didelot for sharing the *B. cereus* dataset. NDM was supported by a James Martin Research Fellowship of the Oxford Martin School. D.J.W. is a Sir Henry Dale Fellow, jointly funded by the Wellcome Trust and the Royal Society (grant 101237/Z/13/Z).

## Supplement

### Algorithm for updating the ancestral material

We record ancestral material of each lineage as a list *m*= (*m*_1_*,…, m_n_*) of pairs *m_i_*= (*s_i_, p_i_*) with *s_i_* integers and *p_i_* real numbers with *p*_0_ = *x_cur_* and *p*_i_ > p_i_−*1*. The first element, *m*1, tells us that from the current position *p*_0_ = *x_cur_* until position *p*_1_, the considered lineage is constantly ancestral to *s*_1_ samples. The second element, *m*_2_, if present, tells us that between positions *p*_1_ and *p*_2_ the considered lineage is constantly ancestral to *s*_2_ samples. Similarly for the other elements of *m*. If for any *i ≤ n* we have *s_i_*= 0 or *s_i_*= *N*, where *N* is the number of samples, then the considered lineage is not in local tree between positions *pi−*1 and *p_i_*. Here we discuss how we update in FastSimBac these ancestral material lists after the local ARG is modified by including a new recombination event and lineage. Updating the lists when *x_cur_* is changed is instead trivial, as it just requires to remove all elements of the lists with *pi ≤ x_cur_*. Also, there is no need to modify the lists after some lineages with no ancestral material are removed from the current local ARG.

The update of the lists is performed by a function addMRCAUp(*l*, *m’*) that iteratively travels upward along the tree starting from lineage *l*, and updates the ancestral material lists of the lineages it encounters by summing ancestral material *m’* to it. addMRCAUp is called twice; the first time it is called on the new recombinant lineage *l*1 with a positive *m’*(the ancestral material of the new lineage has to be added to the lineages above the one it coalesces to). If recombination happens on lineage *l* with ancestral material *m*, then *m’* is defined as the intersection of *m* with the recombining segment *m’*= *m ∩*[*x_cur_, x_end_*] (*m’* has the same values as *m* on [*x_cur_, x_end_*], and is 0 after *x_end_*). addMRCAUp(*l*1, *m’*) is then called. If *l*_2_ is the sister lineage of *l*_1_ (the one with which it shares the recombination point), then addMRCAUp(*l*_2_, *−m’*) is also called, where *−m’* is obtained from *m’* by taking the same positions *p_i_* but opposite counts *−s_i_*.

Each call of addMRCAUp(*l*, *m’*) does the following:

1. Add *m’* to the ancestral material of *l*.
2. If *n_l_*, the node above *l*, is the root of the local ARG, end the iteration.
3. If *n_l_* is a coalescent node with parent lineage *l’*, call addMRCAUp(*l’*, *m’*).
4. Otherwise if *n_l_* is a recombination node with recombinant parent lineage *l*_1_ and sister *l*_2_, and if [*p*_1_*, p*_2_] is the recombining segment of *n_l_*, then we will call addMRCAUp(*l*_1_, *m’ ∩*[*p*_1_*, p*_2_]), and addMRCAUp(*l*_2_, *m’ \*[*p*_1_*, p*_2_]). addMRCAUp(*l*, *m’*) is not executed if *m’* is empty.

### Features of FastSimBac

FastSimbac can be used to generate a sequence of local trees for subsequent intervals of the genome. For example local trees can be written to output with the option “-T”:

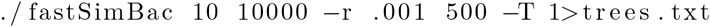

generates a sequence of local trees in the file “trees.txt” for a genome of 1000 bp, 10 isolates, a recombination initiation rate of *ρ*= 0.001, and a mean recombination tract length of *λ*= 500. SNPs can also be generated by specifying a mutation rate with option “-t”, for example:

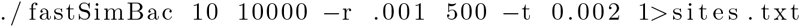

simulates mutations along the local trees (generated similar to before) under a per-base scaled mutation rate of *θ*= 0.002 and under a two-allele infinite sites model, and writes the list of SNPs generated in the file “sites.txt”. For more realistic mutation scenarios, the local trees generated by FastSimBac can be used as input in SeqGen (Rambaut and Grassly, 1997).

Complex evolutionary histories of population structure and demography can be specified using options similar to ms (Hudson, 2002) and MaCS (Chen et al., 2009). For example, *n* populations with migration rate (backward in time) of *m*= 0.1 between them can be specified with the option “-I”:

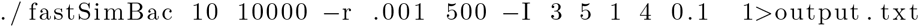

simulates 3 populations (with 5 samples from the first, 1 sample from the second, and 4 samples from the third) with migration rate of 0.1 (the migration rate for each population is 0.1 divided by the number of populations minus one). Population growth can be simulated with option “-G”:

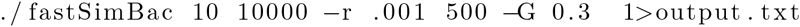

Simulates under an exponentially growing (*α*= 0.3 in this case, or shrinking if *α <*0) effective population size *N*(*t*) = *N*_0_ exp(*−αt*), where *N*0 is the population size at time 0 (present) and *t* is the time before the present, measured in units of 2*N*0 generations. Further simulation options depending on a specific time *t*(such as population splits and merges) are described in the list of options at the end of this section.

We can also simulate recombination between populations or species, similar to (Brown et al., 2015), but with some differences. In fact, instead of modeling a generic diverged donor of recombining segments as (Brown et al., 2015), we instead explicitly model different population/species and migration of recombinant segments between them. This makes simulations more realistic, as we consider the possibility of multiple donor species/populations with a given set of divergence times, and we also model the coalescent process of recombinant segments within a donor species/population. However, this also makes our model of inter-population recombination more computationally demanding. An example of usage is

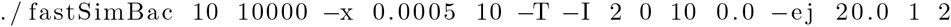

which will simulate 10 samples collected all from the same population, a second unsampled population, no migration between the two populations (these 3 factors are specified by option “-I 2 0 10 0.0” as in ms), a merging backward in time of the two populations at time 20 (option “-ej 20.0 1 2” as in ms), and a cross-population recombination rate of 0.0005 and tract length of 10 (these last 2 features are specified by option “-x 0.0005 10”). So this scenario is useful to simulate two species diverged a certain time in the past but that exchanged small pieces of the genome continuously after being diverged.

Option “-C” can be used to specify an input clonal frame, on which simulations will be conditioned. leaves of the clonal frame must be named with integer numbers from 0 on. An example of usage is:

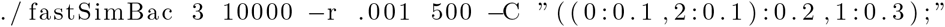

Lastly, option “-R” instructs the program to use variable recombination rate as specified in an input file, and can be used to specify recombination hotspots and coldspots, for example.

The full list of option is:

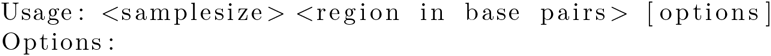

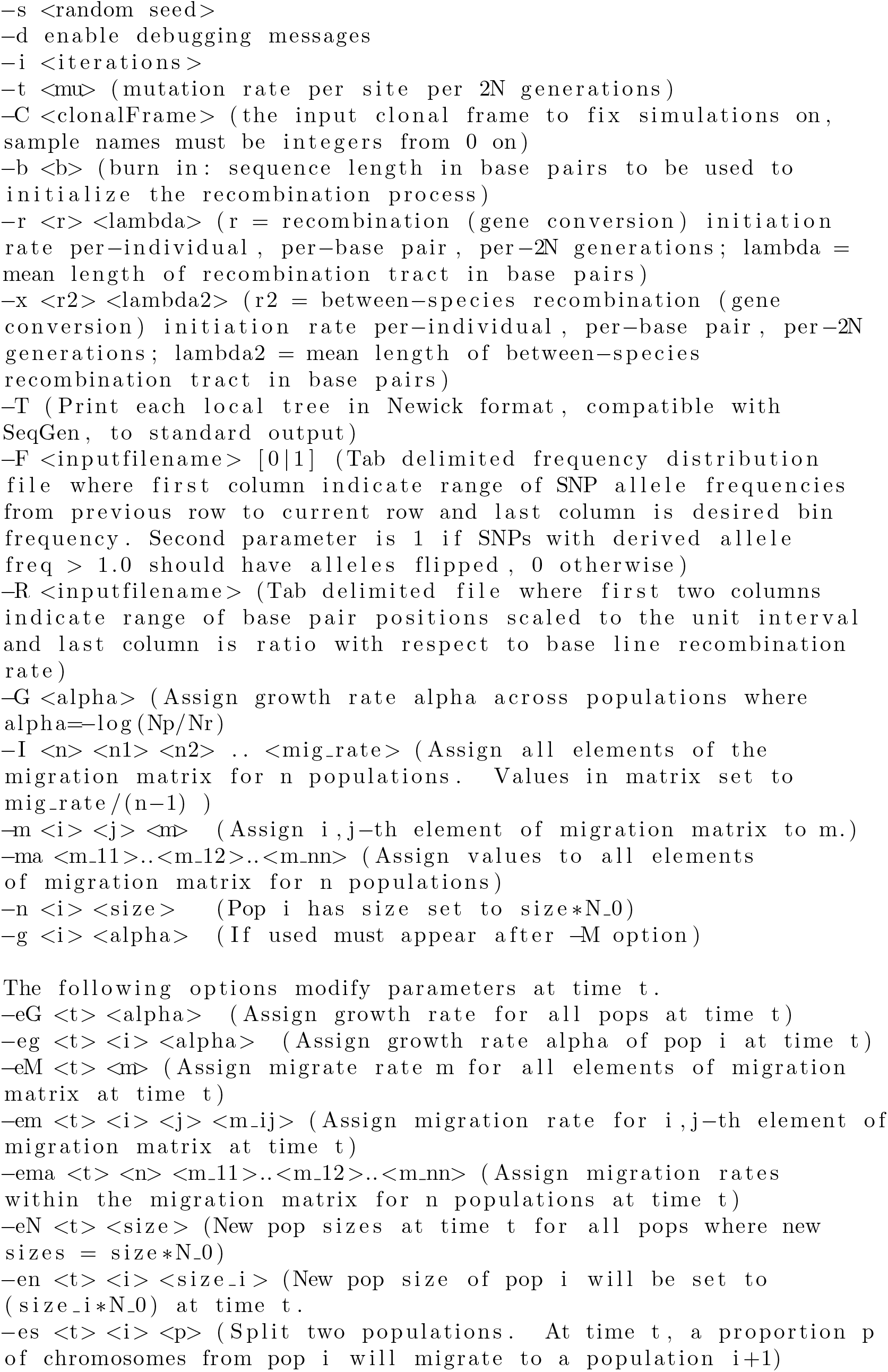

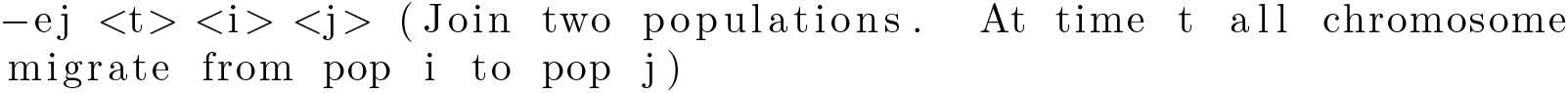

### Correction of Branch Lengths in the ABC-MCMC Analysis

Let us assume that two samples have a time to most recent common ancestor (TMRCA) within the clonal frame of *t*, measured in number of 2*N_e_* generations. Because of recombination, the local TMRCA at any site of the genome between these two samples might be different from *t*. For example, if the two samples are closely related, we suggest that recombination events will likely cause their divergence to increase along some tracts of the genome. On the other hand, if the samples are distantly related, recombination might reduce their phylogenetic distance in some parts of the genome. More precisely, let us consider any given position of the genome, and let us denote with *P*_2_(*x*) the probability that at time *x*(in units of 2*N_e_* generations) the ancestral lineages of the two samples have not coalesced and are both in the clonal frame, with *P*_1_(*x*) the probability that one of them is in the clonal frame and one is recombinant, with *P*_0_(*x*) the probability that none of them is in the clonal frame and so both are recombinant and have not coalesced yet, and finally with *P_c_*(*x*) the probability that at time *x* the lineages have already coalesced. Then, we have by definition that *P_c_*(*x*) = 1 *−*(*P*_0_(*x*) + *P*_1_(*x*) + *P*_2_(*x*)), that *P*_2_(0) = 1 and *P*_0_(0) = *P*_1_(0) = *P_c_*(0) = 0, and that *P*_2_(*x*) = 0 for *x > t*. Let us start by considering values of *x < t*. For these, given that the coalescent rate between any two lineages is 1, the following system of differential equations holds:

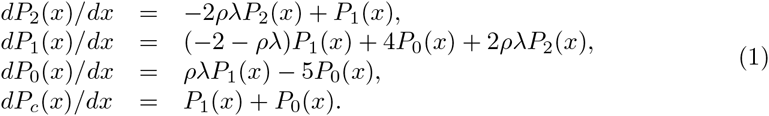

At any value of *x > t*, the expected time to coalesce for the two lineages is 1. So the overall expected time to coalesce for two lineages is:

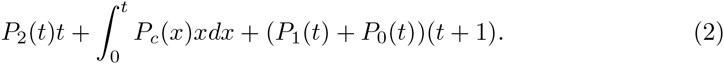

Equation 2 gives us the mean divergence that we expect to observe overall the genome between any two lineages given a clonal frame divergence of *t*. By assuming that the genome is sufficiently large such that the observed divergence corresponds to the expected divergence, inverting this function, we can infer *t*. We performed this numerical integration to infer *t* both in R and Python and verified that we obtained consistent results. We show values of Equation 2 in Figure S1. As can be seen, recombination pushes small values of divergence up, and large values of divergence down, overall homogenizing divergence among samples pairs.

Equation 2 expects, and returns, values expressed in terms of 2*Ne* generations, while what we can measure from data is only divergence in terms of genetic distance. To translate genetic distances into scaled time divergence, we assumed that recombination is strong enough so that the mean divergence time between all lineage pairs is 1, and we ignore recurring mutations.

### Invariant Sites in ABC-MCMC analysis

We get a rough estimate of the transition/transversion rate ratio *κ*= 5.21 as the ratio of the number of observed biallelic SNPs involving a transition over the number of observed biallelic SNPs involving a transversion. We assume that *P*_0_ is the proportion

## Supplementary Figures

**Figure S1.**
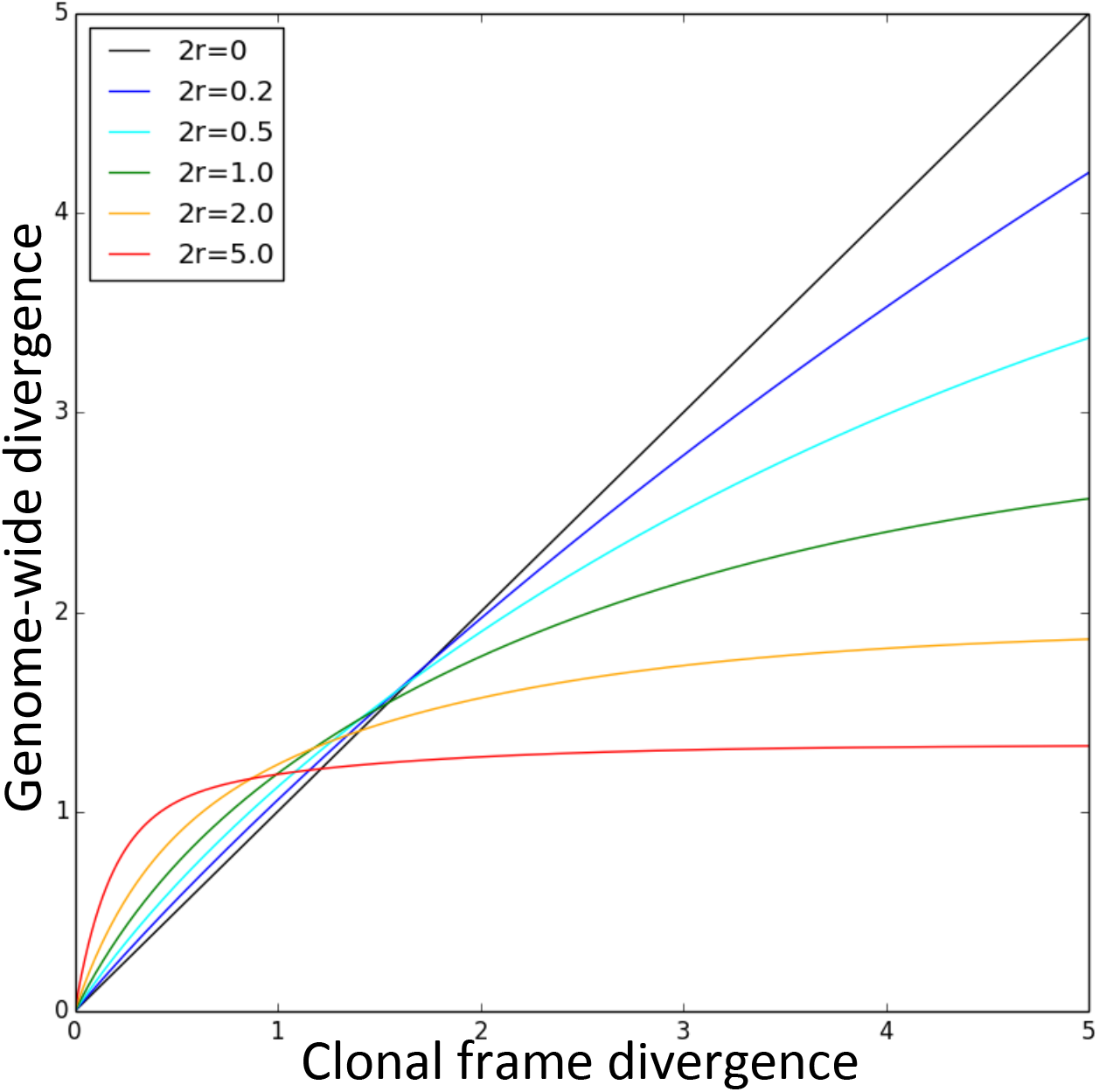
Effect of recombination on mean divergence between pairs of samples. Different lines show the effect of different recombination rates (from *ρ*= 0 to 2*ρ*= 5) as in the legend. On the X axis is the divergence of two lineages in the clonal frame, on the Y axis is the genome-mean divergence after accounting for recombination. of invariant sites, and *μ* the transversion rate times genome-average total tree length. *G*= 3610430 is number of sequenced and aligned sites in the genome. Then we expect about (1 *− P*_0_)*Gμ*(2 + *κ*) biallelic sites and about (1 *− P*_0_)2*μ*^2^*G*(1 + 2*κ*) triallelic sites along the genome alignment. Since we observe 556484 biallelic sites and 73090 triallelic sites, substituting these values in the previous equations, we get a back of the envelope estimate of *P*0 = 0.484. While we are aware that our calculations are very approximate and that the concept of invariant sites itself is an approximation of the more realistic scenario of different degrees of selection affecting different sites, we only use these calculations here to test if accounting for selection can have a strong effect on the inference of recombination parameters.

**Figure S2.**
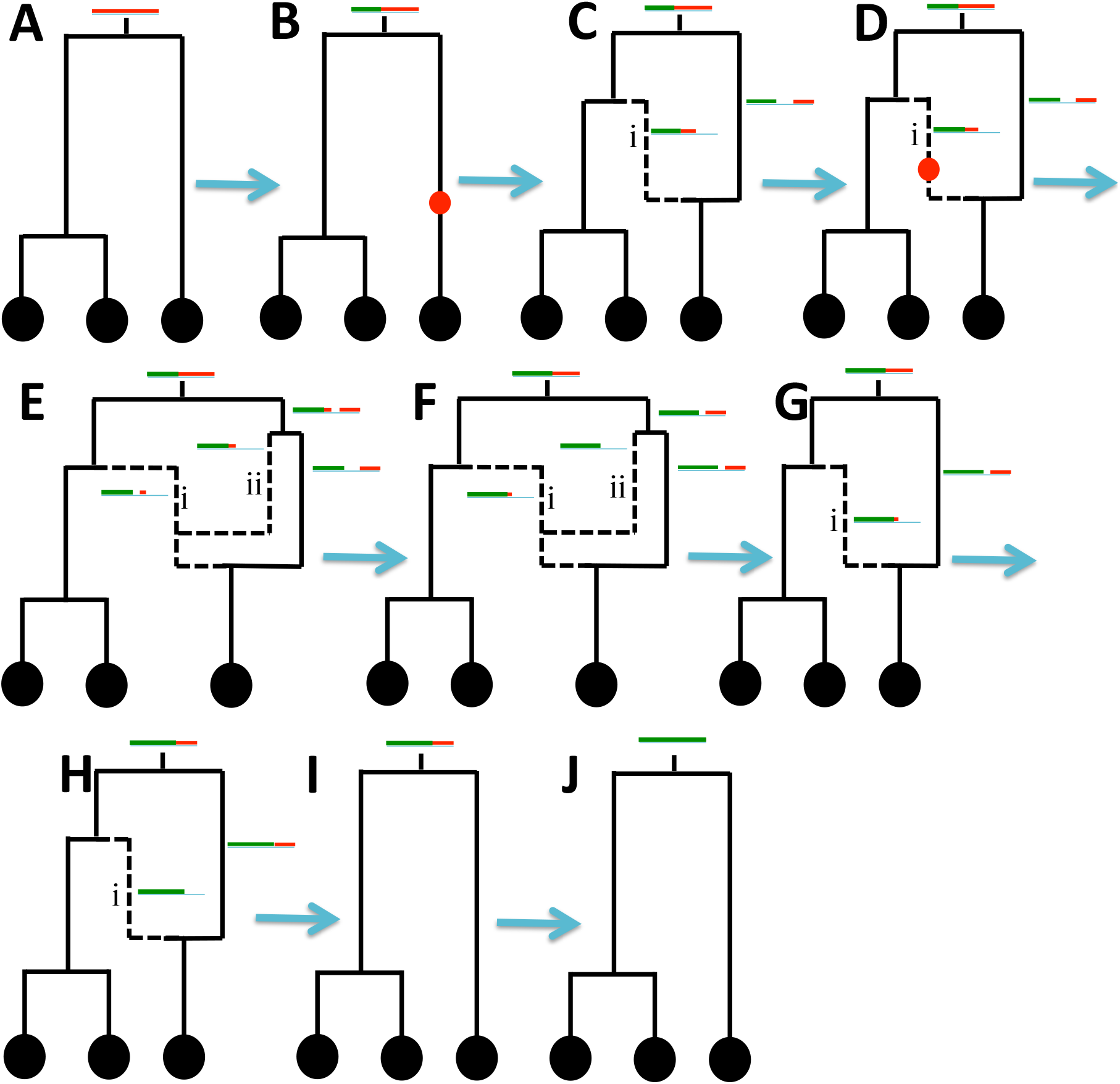
Graphical representation of the BSMC model. Black circles represent samples, black lines are lineages (continuous if they belong to the clonal frame, dashed otherwise). Red segments represent, for each extant lineage, the portion of the genome that is ancestral to any sampled descendent of that extant lineage. Merges of lineages are coalescent events. The process moves left to right across the genome, and left portions of the genome are gradually forgotten (represented in green). **A)** The clonal frame is simulated at the left end (the start) of the genome; **B)** the first recombination event is sampled at the position of the red circle; **C)** the first recombining lineage is created (the dashed line, i) and is coalesced to the rest of the tree; **D)** a second recombination event is sampled (red circle), this time along a recombining lineage; **E)** the second recombining lineage is created (ii), and coalesced to the tree; **F)** the endpoint of the second recombination is reached, the second recombining lineage ii has no ancestral material left; **G)** the second recombining lineage ii is removed; **H)** the endpoint of the first recombination is reached, the recombining lineage i has no ancestral material left; **I)** the first recombining lineage i is removed; **J)** the right end of the genome is reached.

**Figure S3.**
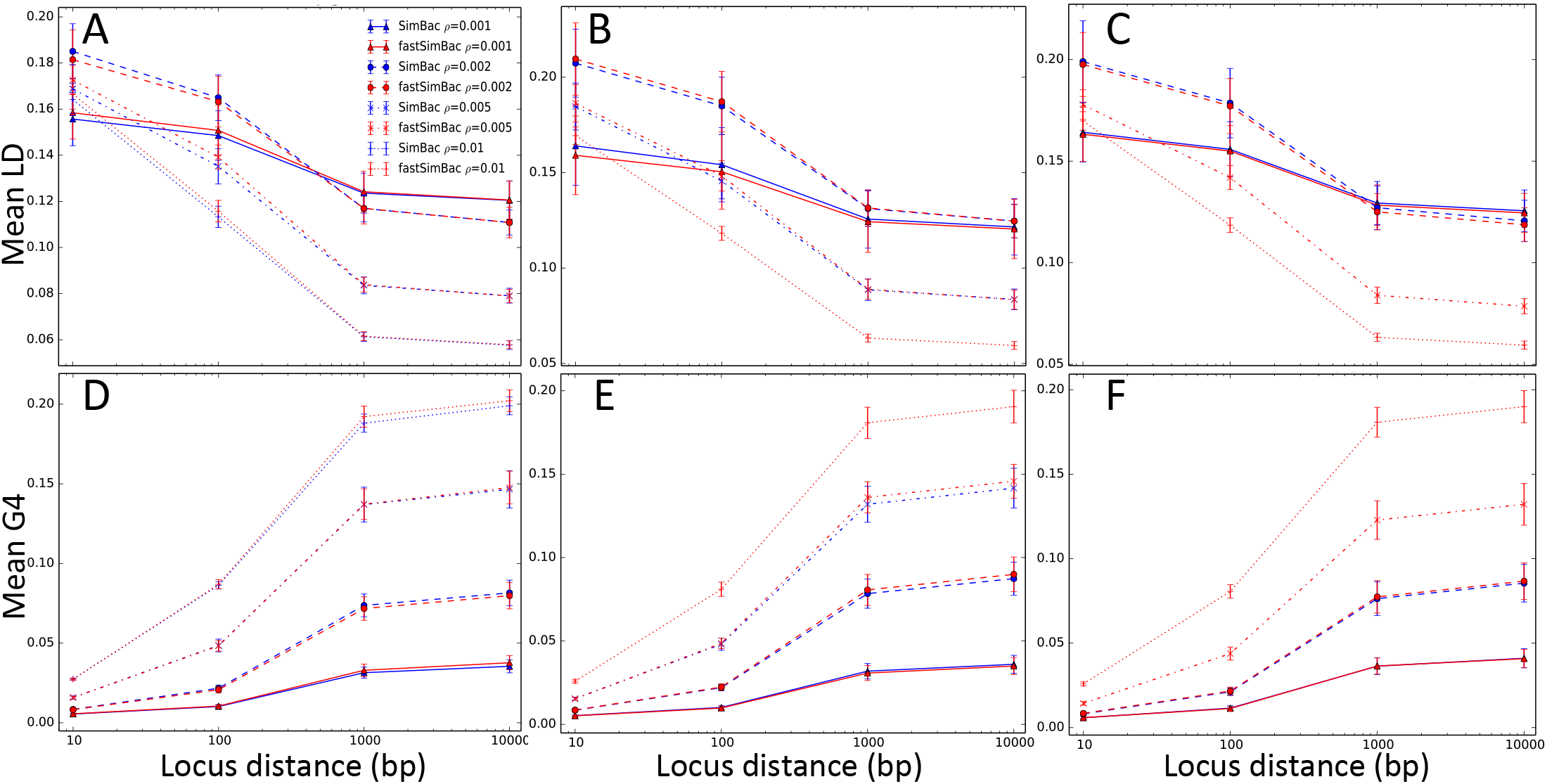
Comparison of linkage disequilibrium and site incompatibility patterns at longer genome sizes. The BSMC simulates patterns of linkage disequilibrium (LD) and site incompatibility (G4) very similar to the bacterial coalescent (or CGC). LD is calculated as 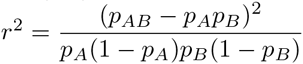. G4 (the four-gamete test) is measured as the proportion of incompatible SNP pairs. On the X axis is the distance (in bp) between SNPs in a pair at which LD and G4 are calculated. For each distance *d* on the X axis, and for any SNP *x* in the alignment, LD and G4 are calculated between *x* and the first SNP at least *d* bp to the right of *x*. Red lines refer to FastSimBac, blue lines to SimBac, and different point and line styles refer to different recombination rates (see legend). Each point is the mean over 20 replicates, and bars are standard errors of the mean. **A)** Mean LD for a genome of 2Mbp, **B)** 5Mbp, **C)** 10Mbp. **D)** Mean G4 for a genome of 2Mbp, **E)** 5Mbp, **F)** 10Mbp. SimBac was not run for the highest recombination rates and genome sizes due to time limitations.

**Figure S4.**
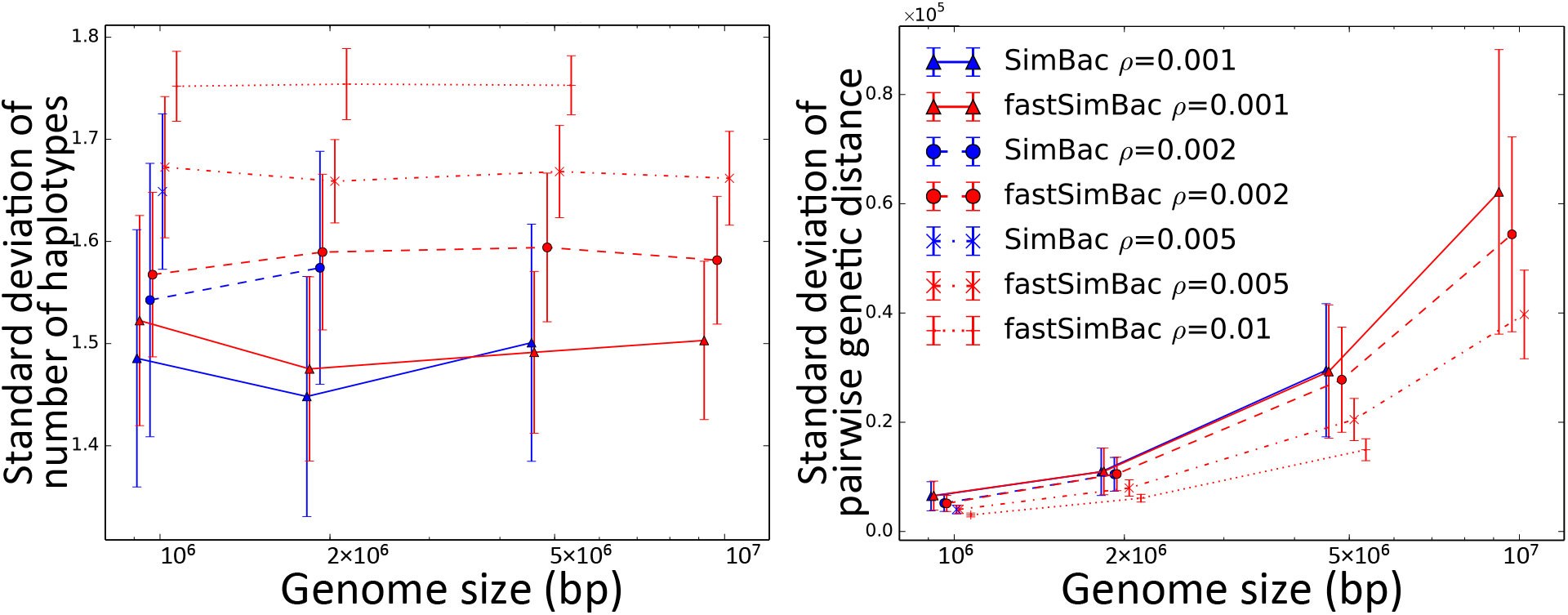
Comparison of level of genome-wide variation in genetic structure simulated between the BSMC and the CGC. The BSMC and the CGC show similar simulated variation in genetic structure along the genome, despite the approximation in the BSMC. **A)** Genome-wide standard deviation of number of simulated haplotypes across non-overlapping sliding windows of 10 SNPs; higher values mean that different loci have more different numbers of haplotypes. **B)** Standard deviation across sample pairs of the number of whole-genome genetic differences; lower values mean that genetic distances among sample pairs are more homogeneous (expected with higher recombination rates). On the X axis the genome size (in bp and on log scale). Red lines refer to FastSimBac, blue lines to SimBac, and different point and line styles refer to different recombination rates (see legend). Each point is the mean over 50 replicates, and bars are standard deviations. SimBac and FastSimBac were not run for the highest recombination rates and genome sizes due to time and memory limitations.

**Figure S5.**
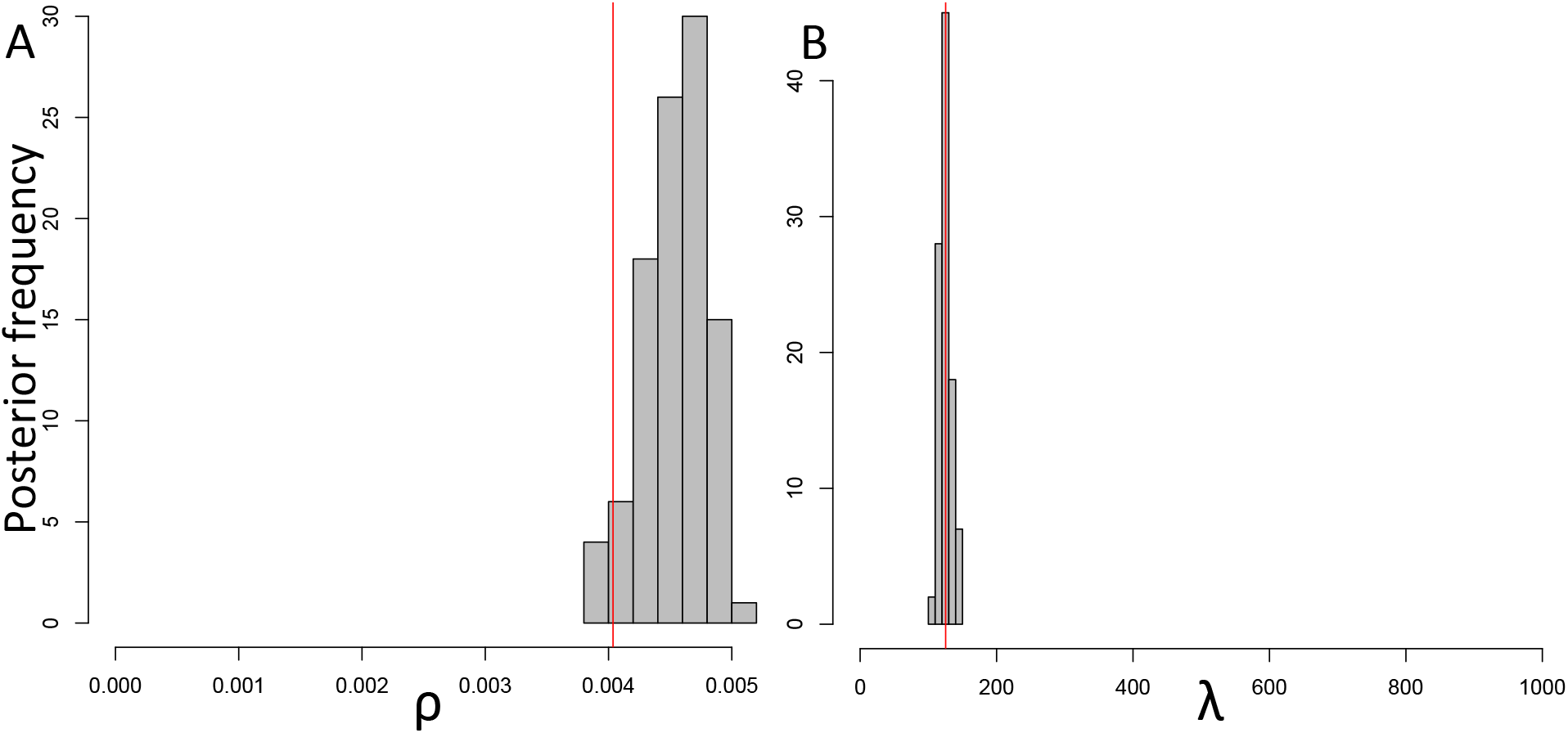
Second replicate of ABC inference using the BSMC on data simulated under the CGC. Recombination parameters simulated under the exact CGC (red vertical lines) where reconstructed using simulations under the BSMC within an ABC inference scheme. **A)** Posterior distribution of *ρ*. **B)** Posterior distribution of *λ*.

**Figure S6.**
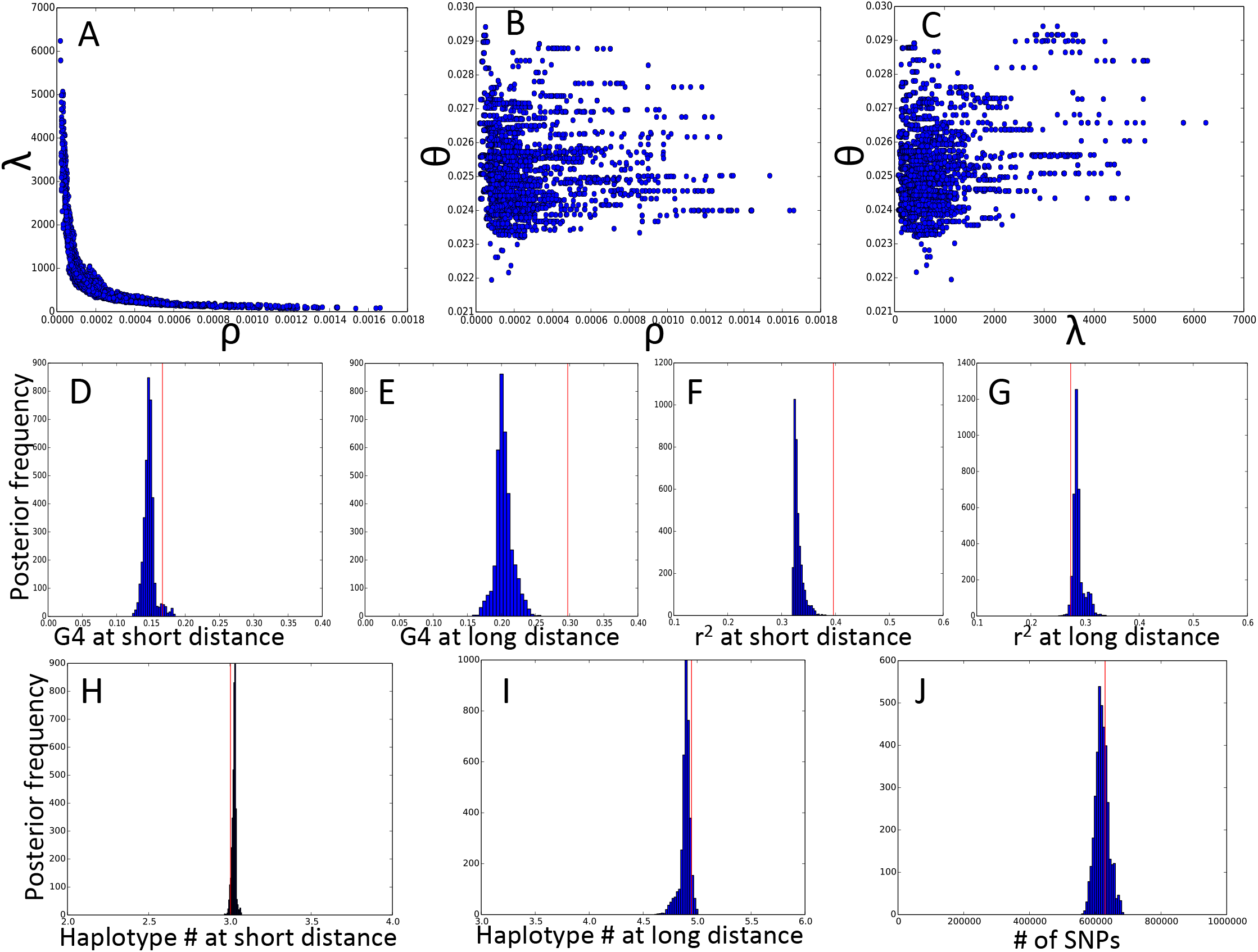
Scatterplots of posterior parameters and summary statistics for genome-wide evolution of *B. cereus*. We inferred BSMC parameters using an ABC-MCMC inference scheme. In the first three scatterplots we show posterior distributions of **A)** *ρ* and *λ*; **B)** *ρ* and *θ*; **C)** *λ* and *θ*. In the second and third rows we show the posterior distributions of summary statistics: **D)** G4 (proportion of SNP pairs that are not consistent, breaking the 4-gamete rule) for consecutive SNPs (“short distance”); **E)** G4 for SNPs at least 2kbp away (“long distance”); **F)** mean linkage disequilibrium (LD, measured as 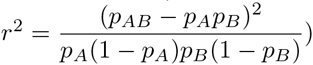 for consecutive SNPs (“short distance”); **G)** mean linkage disequilibrium for SNPs at least 2kbp away (“long distance”); **H)** mean number of haplotypes for pairs of consecutive SNPs (“short distance”); **I)** mean number of haplotypes for groups of 4 SNPs made of 2 pairs of consecutive SNPs, the two pairs being at a distance of at least 2kbp (“long distance”); J) number of SNPs. Summary statistics of the real dataset are shown with red vertical lines in plots **D-J)**.

**Figure S7.**
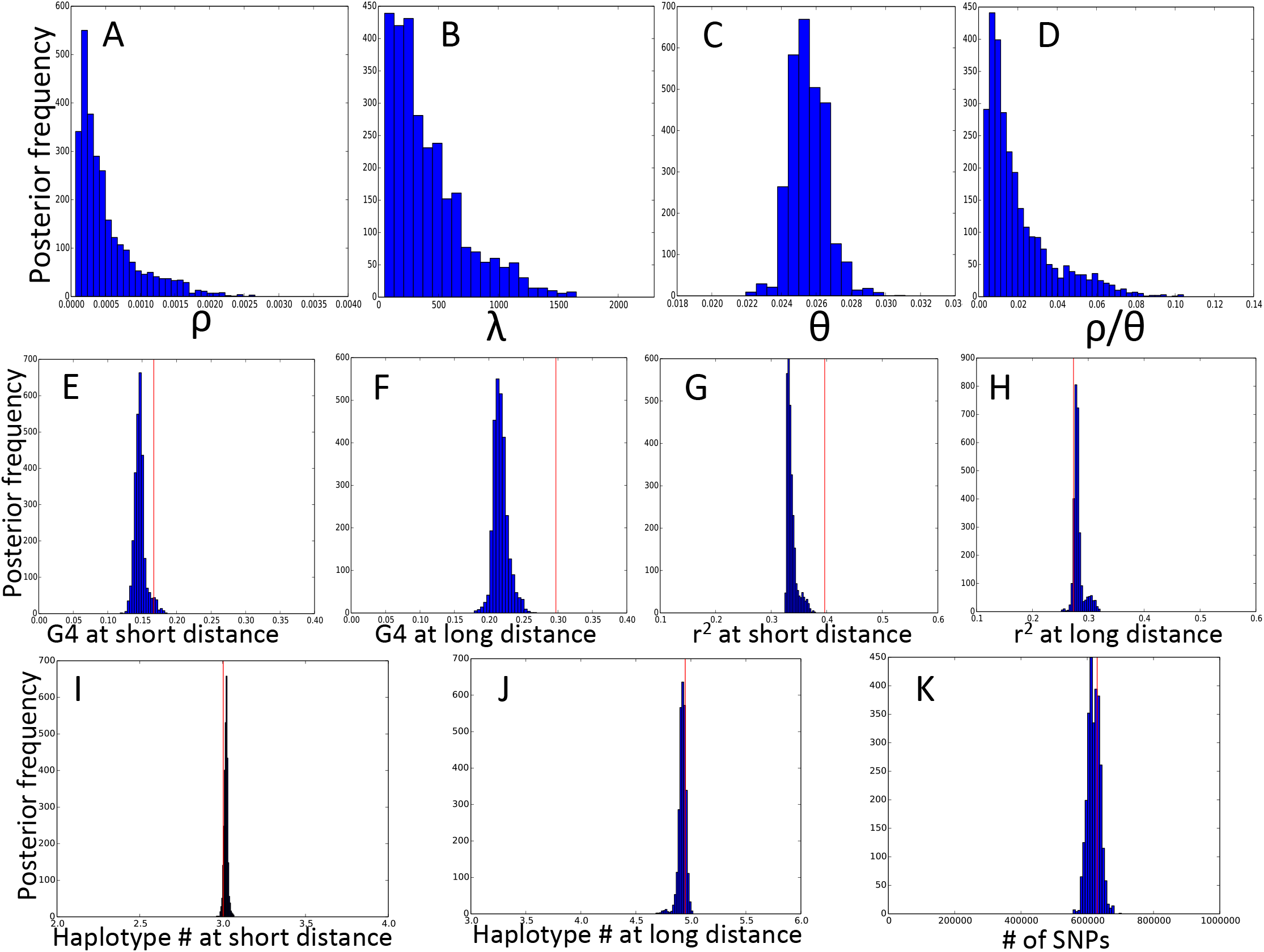
Posterior distributions of parameters and scatterplots of summary statistics for a second inference of genome-wide evolution of *B. cereus*. We inferred BSMC parameters using an ABC-MCMC inference scheme, similar, but independent, from the one in Figures 7 and S6. **A)** Posterior distribution of *ρ*(interquartile range [0.0002,0.0007]). **B)** Posterior distribution of *λ*(interquartile range [181,563]). **C)** Posterior distribution of *θ*(interquartile range [0.0247,0.0262]). **D)** Posterior distribution of *ρ/θ*(interquartile range [0.008,0.026]). **E)** Posterior distributions of G4 (proportion of SNP pairs that are not consistent, breaking the 4-gamete rule) for consecutive SNPs (“short distance”); **F)** Posterior distributions of G4 for SNPs at least 2kbp away (“long distance”); **G)** Posterior distributions of mean linkage disequilibrium (LD, measured as 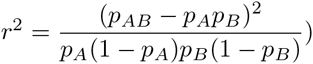) for consecutive SNPs (“short distance”); **H)** Posterior distributions of mean linkage disequilibrium for SNPs at least 2kbp away (“long distance”); **I)** Posterior distributions of mean number of haplotypes for pairs of consecutive SNPs (“short distance”); **J)** Posterior distributions of mean number of haplotypes for groups of 4 SNPs made of 2 pairs of consecutive SNPs, the two pairs being at a distance of at least 2kbp (“long distance”); K) Posterior distributions of number of SNPs. Summary statistics of the real dataset are shown with red vertical lines in plots **E-K)**.

**Figure S8.**
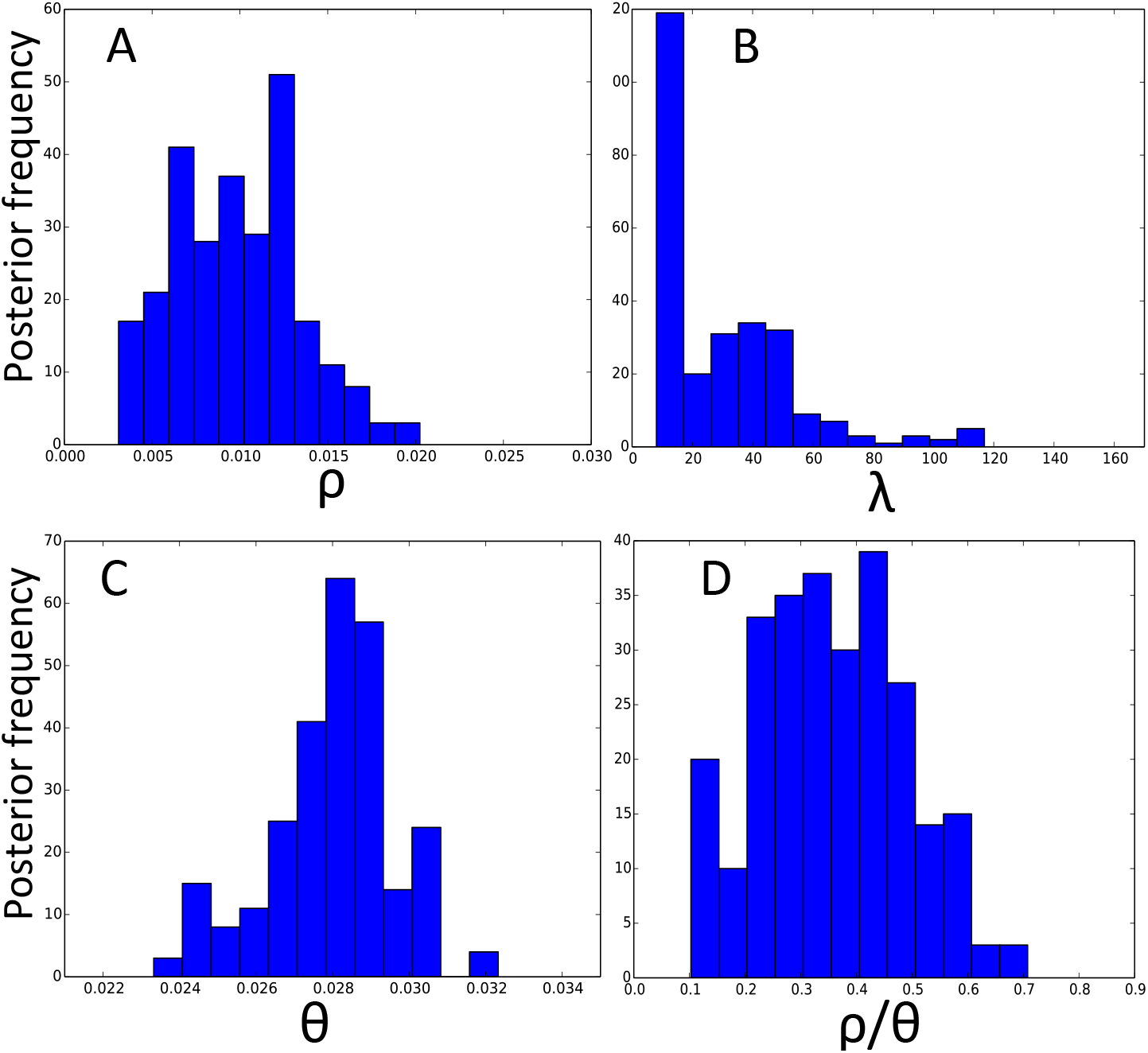
Posterior distributions of parameters for genome-wide evolution of *B. cereus* when not accounting for invariant sites. We inferred BSMC parameters using an ABC-MCMC inference scheme as in Figure 7, but this time without account for invariant sites, without correcting branch lengths, an running for 5000 ABC-MCMC steps. **A)** Posterior distribution of *ρ*(interquartile range [0.007,0.013]). **B)** Posterior distribution of *λ*(interquartile range [14,44]). **C)** Posterior distribution of *θ*(interquartile range [0.0271,0.0289]). **D)** Posterior distribution of *ρ/θ*(interquartile range [0.257,0.449]).

